# Open-source modular FPGA system for two-photon mesoscope enabling multi-layer, multi-depth neural activity recording and lifetime imaging

**DOI:** 10.1101/2025.08.13.669807

**Authors:** Riichiro Hira, Fumiya Imamura, Hiroto Imamura, Yuki Yoneyama, Takehisa Handa, Osamu Fujioka, Che-Hang Yu, Satoshi Suitoh, Reiko Hira, Atsushi Kamoshida, Shigeki Kato, Kazuto Kobayashi, Hiroki Shiwaku, Hidehiko Takahashi, Spencer L. Smith, Akihiro Funamizu, Yoshikazu Isomura

**Affiliations:** Department of Physiology and Cell Biology, Graduate School of Medical and Dental Sciences, Institute of Science Tokyo, Tokyo, 113-8519, Japan; High Performance Artificial Intelligent System Research Team, Center for Computational Science, RIKEN, Saitama 103-0027, Japan; Institute for Quantitative Biosciences, the University of Tokyo, Laboratory of Neural Computation, Tokyo 113-0032, Japan; Department of Life Sciences, Graduate School of Arts and Sciences, the University of Tokyo, Tokyo, 153-8902, Japan; TRIONIX Inc., Saitama 330-0063, Japan; Department of Electrical and Computer Engineering, University of California Santa Barbara, Santa Barbara, CA, 93106-5100, USA; KYOCERA SOC Corporation, Kanagawa 226-0006 Japan; Graphical Design Lab, Saitama 330-0843, Japan; Department of Molecular Genetics, Institute of Biomedical Sciences, Fukushima Medical University, Fukushima, 960-1295, Japan; Department of Psychiatry and Behavioral Sciences, Tokyo Medical and Dental University Graduate School, Tokyo 113-8510, Japan

**Keywords:** FPGA, 4 plane imaging, two-photon, multiplexing, mesoscope, lifetime imaging, open-source, cerebral cortex

## Abstract

Large field-of-view (FOV) two-photon microscopy makes it possible to record a large number of neural activities from multiple brain regions simultaneously. However, the larger the field of view, the longer it takes to scan the entire FOV. To increase imaging speed, we have developed open-source software to digitize analogue signals from a photomultiplier tube using a field-programmable gate array (FPGA) at a rate of 3.2 GS/s. By combining this with a newly developed a circular delay-path module for a custom two-photon mesoscope (Diesel2p), we succeeded in simultaneous recording of >10,000 neurons from the entire bilateral dorsal cortex at up to four depths. We also demonstrated large FOV lifetime imaging using the same system. Our modular, open-source FPGA system can be readily integrated into any type of two-photon microscope and will accelerate the biomedical application of multi-scale two-photon imaging in a wide range of pathophysiological investigations.

## Introduction

Large field-of-view (FOV) two-photon microscopy has advanced systems neuroscience by enabling simultaneous recording of thousands of neurons across multiple brain regions (Tsai et al. 2015; Yu et al. 2024, 2021; Clough et al. 2021; Stirman et al. 2016). Two-photon microscopy requires the laser to be focused on a single point in space and time; therefore, a wider FOV leads to longer scan durations. Imaging speed can be improved either by faster point scanning or by multifocal imaging. Acousto-optic deflectors (AODs) (Duemani Reddy et al. 2008; Grewe et al. 2010), tomographic methods (Kazemipour et al. 2019), and temporal focusing (Oron, Tal, and Silberberg 2005; Zhu et al. 2005; Prevedel et al. 2016; Weisenburger et al. 2019) can also increase the sampling rate, but remain challenging to implement over large FOVs. Multifocal excitation using a beam splitter (Nielsen et al. 2001), a spatial light modulator (SLM) (Onda et al. 2021; Han, Yang, and Yuste 2019), or a spinning-disk (Maeda et al. 2018; Otomo et al. 2020) can raise the frame rate. A delay-line strategy combined with high-speed sampling can further accelerate acquisition by several-fold and is compatible with large FOV two-photon imaging (A. Cheng et al. 2011; Yu et al. 2021; Stirman et al. 2016; Amir et al. 2007). Extensions of this approach have achieved even faster scanning (Lai et al. 2021; Beaulieu et al. 2020; Demas et al. 2021; Weisenburger et al. 2019). Delay line strategies usually rely on low-repetition-rate lasers because the longer interval between pulses allows distinct temporal delays to be inserted for efficient multiplexing. Introducing such delays is a key route to faster imaging in large FOV two-photon microscopes. However, high-speed sampling in this context requires detailed knowledge of the detection electronics, and high-power low repetition lasers remain uncommon. An add-on module that uses the standard 80 MHz laser in existing systems to create a multifocal source would therefore markedly increase throughput and accelerate progress in systems neuroscience.

In addition to the fluorescence intensity, a two-photon microscope can measure fluorescence lifetimes by comparing the arrival time of fluorescence photons with that of the excitation pulses (Torrado et al. 2024; Datta et al. 2020; Park and Gao 2024). There are two main approaches. Time-correlated single-photon counting (TCSPC), which provides the highest temporal resolution (Yasuda et al. 2006; Gratton et al. 2003; Murakoshi, Wang, and Yasuda 2011), and digitizing the photomultiplier-tube (PMT) signal at high sampling rates to reconstruct decay curves digitally (Eibl et al. 2017; Bower et al. 2018; Dow et al. 2015; S. Cheng et al. 2014; Giacomelli et al. 2015; Sorrells et al. 2021). TCSPC excels when photon counts are low, whereas high-speed digitization is more scan-efficient for bright specimens that produce many photons, although it requires rapid digital processing. Open-source software that can handle these high-throughput data streams would bring significant benefits not only to neuroscience but also to biology and medicine.

In this study we developed a laser multiplexing mechanism and an open-source software module for high-speed sampling, then integrated them with the Diesel2p mesoscope (Yu et al. 2021; Hira et al. 2023; Clough et al. 2021). The module samples the PMT signal at 3.2 GS/s without dead time, and at each 80 MHz laser pulse the FPGA computes fluorescence values from four independent focal points in parallel. With this setup we imaged neural activity in a mouse brain at four focal planes simultaneously and also demonstrated large field-of-view two-photon lifetime imaging using the same system. Because the fast analog sampling device is modular and can be added to any two-photon microscope, it promises broad benefits across biomedical research.

## Materials and methods

### Animal preparation

All experiments were approved by the Institute of Science Tokyo (A2021-290C6, A2024-060C2) and were carried out in accordance with the Fundamental Guidelines for Proper Conduct of Animal Experiment and Related Activities in Academic Research Institutions (Ministry of Education, Culture, Sports, Science and Technology, Japan). We used adult male and female double-transgenic mice (AI162 or TIT2L-GC6s-ICL-tTA2 (Daigle et al. 2018) × Slc17a7-IRES2-Cre (Madisen et al. 2015)) and wild-type mice. These mice were kept under an inverted 12h/12h light-dark schedule (lights off at 12:00 AM; lights on at 12:00 PM) in their home cages to match experimental timing.

### Neonatal AAV injection

Neonatal viral injection was done on postnatal day (P)1-2 (Oomoto et al. 2021). The pups were anesthetized by hypothermia on an ice bed. A quartz glass pipette (outer diameter 50 µm) was inserted into the cortex to a depth of 250–300 µm below the pial surface. A total of 4 µL of AAV-DJ-Syn-jGCaMP8s (Zhang et al. 2023) was delivered over 1 min.

### Surgery

Craniotomy over the dorsal cerebral cortex was performed under isoflurane anesthesia (1.5– 1.8%). Body temperature was maintained using air-activated heat packs, and corneas were protected with ophthalmic ointment. The scalp overlying the dorsal cortex was removed, and a custom head-fixation imaging chamber was mounted to the skull with a self-curing dental adhesive (Super-Bond, L-Type Radiopaque; Sun Medical) and luting cement (Fuji Lute). The surface of the exposed skull was subsequently coated with clear acrylic dental resin (Super-Bond, Polymer Clear; Sun Medical) to prevent drying. Mice were then returned to their home cages after recovery.

### Cranial window implant

Three days after head-plate surgery, we performed a large cranial window implant. The glass plug used to cover the dorsal cortex was made with a 6 mm × 6 mm, #3 coverslip and a 7 mm × 7 mm, #0 coverslip with two corners cut (Matsunami Glass). The two coverslips were bonded using optical adhesive (Norland Optical Adhesive 63; Norland Products). After anesthesia, 20% mannitol (5 µL per g body weight, intraperitoneal) was administered to reduce intracranial pressure, and atropine sulfate (0.216 µg per g, intraperitoneal) to reduce saliva and mucus secretion. The head plate was secured in a stereotaxic frame. Previously applied acrylic resin was removed using a dental drill, followed by marking of bregma and stereotaxic coordinates. The skull along the outline of the cranial window as well as the coronal and sagittal sutures was thinned with the drill. The frontal and parietal bones were lifted in four separate pieces. The glass plug was then placed on the exposed cortex, enabling optical access to the entire dorsal cortex, and sealed to the bone with cyanoacrylate glue and dental resin cement. Finally, the mouse was administered mannitol (10 µL per g, intraperitoneal) and meloxicam (0.38 µL per g, subcutaneous) before returning to the cage to recover for five days.

### Two-photon imaging

All two-photon imaging was conducted with a custom-made mesoscope, Diesel2p (Yu et al. 2021). For excitation, we used a fixed-wavelength fiber laser, ALCOR 920-2 (Spark Lasers, France). The group-delay dispersion (GDD) compensation, determined by maximizing the fluorescence intensity of a sample (≈ 60,000 fs²), was adjusted within the laser. A 2× or 3× beam expander (GBE02-B, GBE03-B, Thorlabs) expanded the laser to fill the back aperture of the objective lens. Two independent scan engines (Path-1 and Path-2), each comprising two galvanometer mirrors (R6220H, Cambridge Technology; GS7X-AG, Thorlabs) and an 8 kHz resonant scanner (CRS8kHz, Cambridge Technology), together with an electrically tunable lens (ETL, EL-12-30-TC-VIS-16D; Optotune) and a shutter (SHB05T, Thorlabs), were controlled with a PCI-6722 (National Instruments). The scanner power supplies were custom-made. The PMT (H10770PA-40, Hamamatsu) signal was amplified with either an amplifier (cutoff frequency 200 MHz, gain 100; Edmund Optics #59-179) or a C5594 preamplifier (cutoff frequency 1.5 GHz, gain 36 dB; Hamamatsu Photonics). When using the C5594, the signal was passed through a BNC high-pass filter (EF515; Thorlabs) and attenuated with a 10 dB BNC attenuator (Tyclon, BA-PJ-10) to fit the digitizer’s dynamic range. The signal was digitized with ATS-460 (200 MS/s; AlazarTech) or PXIe-5774 (3.2 GS/s; National Instruments). The synchronization signals from the resonant scanner and the galvo scanner were recorded at 200 MS/s. The PXIe-5774 digitizer was installed in a PXIe-1082 chassis, and the signals were transferred to the host PC via a PXIe-8301 Thunderbolt interface. The entire system was controlled by custom software written in LabVIEW 2019 (National Instruments).

Calibration of imaging scale was carried out empirically with a fluorescent scale. The scale was made by laminating a fluorescent reference slide (FSK5; Thorlabs) to a resolution test target (R1L3S1P; Thorlabs) and fixing the assembly in a slide holder (MAX3SLH; Thorlabs). A goniometric stage with a rotation center height of 125 mm (Misumi, B58-60TLC) and a rotation stage (XXR1/M, Thorlabs) under the objective lens were used to adjust the fluorescent scale until it was parallel to the imaging plane, after which calibration images were acquired.

### Excitation point spread function (PSF) measurements

The measurement and analysis procedures were described in detail previously (Yu et al. 2021). To evaluate the excitation point-spread function (PSF), sub-micrometer fluorescent beads (0.2 µm-diameter, Invitrogen F-8811) were embedded in a ∼1.2 mm-thick 0.75% agarose gel and the gel was sealed with a cover glass (Matsunami, No. 1, 0.12–0.17 mm). Z-stacks ranging from 30 to 100 µm were acquired. The stage was moved axially in 0.2 µm increments (Thorlabs, KVS30). At each focal plane, 10 frames were acquired and averaged to yield a high signal-to-noise image. Due to the difference between the refractive index of the objective immersion medium (air) and the specimen medium (water), the actual focal position within the specimen shifted by Δfocus = 1.35 × Δstage.

### Multiplexing

The pulsed laser for excitation operated at 80 MHz (interpulse interval 12.5 ns). In a circular delay-path module (**CDP module**), an s-polarized laser was reflected by a PBS and routed through an additional 1.88 m loop (c × 6.25 ns ≈ 1.88 m). This produced a second pulse delayed by 6.25 ns. The CDP module contained an electrically tunable lens (ETL) to adjust focal depth. After recombination, the beam passed through a half-wave plate (HWP) and was split again by a second PBS. The beam propagated along either Path-1 or Path-2, with Path-2 being 0.94 m longer, effectively dividing each source pulse into four pulses equally spaced by 3.125 ns (effective repetition rate 320 MHz). The two lateral paths enabled imaging at two different locations simultaneously. Furthermore, the presence of the ETL in the CDP module and achromatic/collection lenses in Path-2 made it possible to scan four planes at different areas and depths. The emitted fluorescence passed through the objective and a dichroic mirror and was detected by a PMT. The signals were then amplified and demultiplexed by an FPGA-based high-speed sampling module. Detailed instructions are provided in the **Supplementary Information**.

Power distribution among the four fields was adjusted as follows. Two sets of HWPs and polarizing beam splitters (PBSs) define two independent ratios: the ratio between the direct and indirect beams and the ratio between the two direct beams (*path* 1_*direct*_ and *path* 2_*direct*_). The ratio between the delayed beams (*path* 1_*indirect*_ and *path* 2_*indirect*_) is then fixed by the latter. Let the direct-to-indirect ratio be 1 : *x* and the ratio of the two direct beams be 1 : *y*. The indirect beams are therefore in the inverse ratio *y* : 1. For an incident power *M*, the powers are

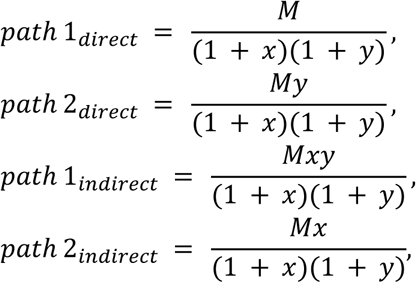

giving the ratio

*path* 1_*direct*_ : *path* 1_*indirect*_ : *path* 2_*direct*_ : *path* 2_*indirect*_ = 1 : *xy* : *y* : *x*. (Here D = direct, I = indirect.)

### Simulation of optical pathways

For each of the four branched optical paths, we optimized the system in Zemax OpticStudio (v16) to ensure that the Strehl ratio exceeded 0.9 at both the field center and at a point 2.5 mm off-axis. The first and second direct paths were configured to allow a coherent laser beam to be incident on the resonant scanner, as in conventional scanning. In contrast, the first and second indirect paths were directed into the resonant scanner via a loop circuit. An electrically tunable lens (ETL, Optotune) was incorporated into the loop circuit, thereby enabling the indirect paths to be scanned simultaneously in the axial direction. Starting from a 4f optical layout, we optimized the parameters by varying the distance between lenses to maintain a Strehl ratio above 0.8 even when the ETL current was altered. During optimization, the light source was set at 920 nm and the beam diameter at the scanner was fixed at 3.6 mm. Although this diameter underfilled the resonant scanner’s 4 mm aperture, it was chosen to ensure sufficiently high laser transmission. The effective numerical aperture was determined via ray tracing. For each of the four optical-path configurations, the focal depth was determined by optimizing the spot radius while varying the distance from the glass window to the focus.

### FPGA-based software

We configured five LabVIEW libraries to control data acquisition, storage, reconstruction, and export of imaging data. The **“Diesel2p Acquisition”** library provides APIs to control the PXIe-5774 FlexRIO digitizer (with on-board FPGA) and to acquire PMT waveforms. The **“TDMS Async Write-Analog”** library streams analog data to an SSD at high speed, and the **“TDMS Async Write-H-sync”** library saves H-sync timing data. The **“Diesel2p File Read”** library reconstructs and displays images from the stored analog and H-sync streams, and the **“Diesel2p File Conversion”** library batch-converts reconstructed images to TIFF files.

Using these libraries, we built four top-level VIs. **“tc-All Data Path for Display Data.vi”** renders images online via the Diesel2p Acquisition API. **“All Data Path for Logging.vi”** saves analog and digital data in TDMS format using the Acquisition API together with both TDMS Async Write libraries. **“Data Viewer.vi”** plays back stored data as video via the File Read API. **“File Conversion for Diesel2p.vi”** exports images from stored TDMS data as TIFF. Each API is encapsulated as a subVI so that components can be reused across projects. Detailed VI and subVI structures are described in the Supplementary Information, and parameter descriptions are available through each subVI’s “Context Help.”

### Real-time data storage and image reconstruction

Data were saved in Technical Data Management Streaming (TDMS) format, a binary container designed for high-throughput streaming in LabVIEW. We stored FPGA-integrated fluorescence counts, the frame-start (V-sync) signal, and resonant-scanner sync (H-sync) signals in TDMS files. Offline reconstruction software reads these TDMS files, selects the scanner sync and channel (1–4), and writes per-path images to binary files. The binary data are then sine-wave corrected to compensate for the position-dependent scan speed of the resonant scanner, and exported as 16-bit TIFF images. These images can be processed with MATLAB, ImageJ, or Python for registration and region-of-interest (ROI) detection, as in conventional two-photon workflows.

### Image processing

Image preprocessing was done using ImageJ (National Institutes of Health) and MATLAB (MathWorks, Natick, MA, USA) software. The bidirectional phase offset of the image was calculated, and sinusoidal distortions caused by the resonant scanner were corrected. The Turgoreg plugin (Thevenaz, et al., 1988) on ImageJ was used for image registration. The registered images were analyzed using Suite2p (Pachitariu et al. 2016) to determine the location of the regions of interest (ROIs). The intensity of each ROI was calculated by subtracting mean values of background pixels within 60 μm of the ROI excluding the other ROIs from mean value of the ROI. For each ROI, the 8 percentile pixel values within 15 s of the time window were obtained, which was then smoothed using a Gaussian filter with standard deviation of 30 s. This was considered as the baseline of neural activity and was subtracted from the original data to set the baseline to 0.

### Single-photon signal measurement

Full-speed data were recorded at 3.2 GS/s in the on-board memory of a PXIe-5774 digitizer, allowing individual photon signals to be resolved. Because the memory capacity is limited, each acquisition lasted ≈ 200 ms. Each PMT trace was processed by applying a threshold and aligning the waveform at the downward crossing point, which we interpret as a single-photon or multi-photon event. Urea crystals were used as the sample. Apart from the sample and laser power, all electronic settings were identical to those used in the in vivo and fluorescence-lifetime imaging experiments.

### Impulse response function (IRF) and fluorescence lifetime imaging

We used Convallaria (standard FLIM specimen). The 80 MHz excitation pulses are separated by 12.5 ns, which the high-speed digitizer divides into 40 temporal bins (Δt = 0.3125 ns). Because the digitizer can store data from only four windows at a time, we acquired ten sequential images with different window positions and concatenated them to create a 40-frame lifetime stack with a temporal resolution of 0.3125 ns per frame. For each pixel this procedure yielded a vector, *y* = [*y*₀, *y*₁, …, *y*₃₉]. To obtain the impulse-response function (IRF) we recorded the second-harmonic signal from a urea-crystal slide, giving, *u* = [*u*₀, *u*₁, …, *u*₃₉]. For convolution we formed a periodic extension of the IRF, *ū* = [*u u u*], to avoid wrap-around at the 12.5 ns boundary.

The fluorescence decay was modeled as the convolution of the IRF with a mono-exponential plus offset, sampled at Δt:

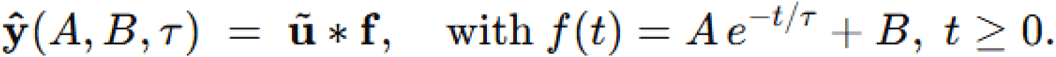

We estimated AAA, BBB, and *τ* by least-squares:

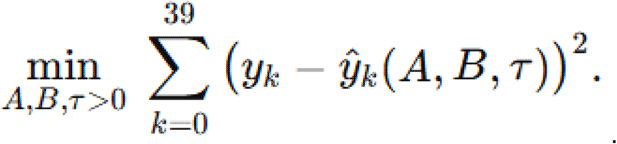

Only pixels brighter than a preset threshold were fitted, and the fitted *τ* values were rendered as the fluorescence-lifetime image.

### Phasor analysis

Phasor analysis was performed as follows. The impulse response function (IRF, 40 bins spanning 12.5 ns) was first normalized to the zero–one range. With the laser repetition period *T* = 12.5 *ns*, the angular frequency was defined as *ω* = 2*π* ⁄ *T*, and the phase of each bin *k* from zero to thirty-nine was set to *θ*_*k*_ = 2*πk* ⁄ 40. The Fourier coefficients of the IRF were then obtained as

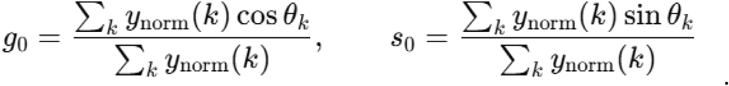

The fluorescence decay trace of every image pixel was likewise normalized to the zero–one range, and its coefficients were calculated as

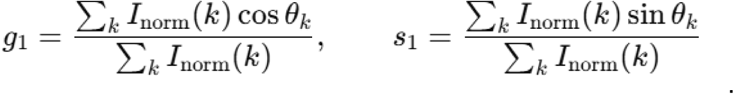

Instrument response was corrected by treating the IRF coefficient *C*₀ = *g*₀ + *i s*₀ and the pixel coefficient *C*₁ = *g*₁ + *i s*₁ as complex numbers and computing *C* = *C*₁ ⁄ *C*₀, which gives *g* and *s* after separation into real and imaginary parts. The phase φ was obtained from arctan (*s* ⁄ *g*), and the lifetime was calculated as

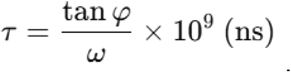

All calculations were executed automatically with a custom MATLAB R2024a script.

### Optimization of detection windows by genetic algorithm

A genetic algorithm was used to choose four temporal accumulation windows from each forty-bin decay waveform (total length twelve point five nanoseconds) so as to maximize the accuracy of single-exponential lifetime estimation with the four-gate method. Before optimization, each specimen and IRF waveform was **circularly shifted by 19 bins** to place the **reference bin at bin 0 (t = 0)**, a **constant baseline was subtracted**, and the result was stored as a **40-sample vector** with **0.3125-ns** spacing. Each gene consisted of eight ordered bin numbers *b*₁ to *b*₈, interpreted as four windows (*b*₁, *b*₂), (*b*₃, *b*₄), (*b*₅, *b*₆), and (*b*₇, *b*₈). The time *t*_*g*_ for a window was taken as the average of its two bin centers multiplied by 0.3125 ns. The observed intensities *Y*_*g*_ for the specimen and *R*(*τ*) for the IRF were defined as the mean intensity in the corresponding bins. Lifetime *τ* for a given gene was obtained by fitting the model

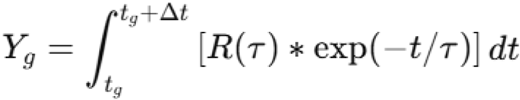

and finding the value of *τ* that minimized the residual sum of squares

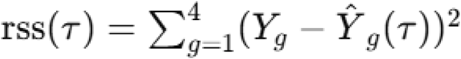

within the interval 0.3125 ns, using a one-dimensional search (*fminbnd*) with a tolerance of one times ten to the minus four. The ground-truth lifetime *τ*_*truth* for fitness evaluation was the 40-bin single-pixel full fit obtained with the conventional method, and fitness for each gene was defined as the Pearson correlation coefficient between *τ* and *τ*_*truth* across 120 randomly chosen pixels. The initial population contained 100 genes and the population evolved for 100 generations. In each generation, the twenty genes with the highest correlation were retained unchanged as elites, and the remaining eighty were produced by selecting an elite at random and shifting one of its bin indices by plus or minus one. Duplicate genes were removed to maintain diversity. The four-window configuration that achieved the highest correlation in the final generation was adopted for subsequent experiments. All computations were carried out automatically with custom MATLAB R2024a scripts.

### Correction of fluorescence leakages

To estimate fluorescence leakage between channels, we imaged a brain slice expressing GCaMP with multiplexing disabled (single 80 MHz beam). and sampled the fluorescence every 0.3125 ns (40 bins per 12.5 ns period). Because the system records four channels in parallel, we acquired 100 frames per setting (four consecutive bins per acquisition) and averaged them, then repeated this procedure while stepping the acquisition window across the 40 bins. This yielded 40 mean images whose pixel intensities reflect the delay of the GCaMP6s signal relative to each laser pulse, given the combined response of the PMT, pre-amplifier, and A/D converter used under the same settings as in the in vivo and fluorescence-lifetime experiments. These temporal profiles were then employed to quantify leakage when four fields of view were imaged concurrently. For each laser pulse the raw pixel values in the four channels were written as *J* = [*J*1, *J*2, *J*3, *J*4]⊤ and the true values in the absence of leakage as *I* = [*I*1, *I*2, *I*3, *I*4]⊤. We assumed a linear mixing model so that *J* = *WI* where *W* is a 4×4 constant matrix whose off-diagonal elements represent cross-talk coefficients determined from the time-resolved calibration data,

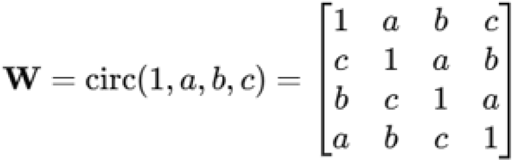

where a, b and c are the leakage coefficients for signals that arrive one, two and three steps earlier, respectively. Applying *I* = *W*^−1^*J* with this explicit inverse removes the leakage among the four channels while preserving the original signal amplitudes. After restoring the pixel values for each individual laser pulse, we reconstructed the images using the start trigger of each resonant scanner sweep. Because our system employs two independent resonant scanners, pixels acquired along the two optical paths do not correspond one-to-one in the raw data. Consequently, instead of correcting the leakage on the reconstructed images, we applied the channel-mixing correction on a pulse-by-pulse basis and then rebuilt the images from the corrected pulse data.

### Statistics

Wilcoxon’s rank sum test was used for pairwise comparison. Data was expressed as means ± S.E.M unless otherwise noted.

## Results

### Laser-efficient four-way beam splitting

We designed optical pathways in which the 80 MHz laser with an inter-pulse interval of 12.5 ns is divided into four parts, with one pulse reaching the specimen every 3.125 ns. First, a loop optical path (1.88 m per loop) was constructed so that the s-polarized laser reflected by the PBS makes one loop. The electrically tunable lens (ETL) was placed in the loop for focal-depth adjustment under the objective. Two concave lenses (f=500) were added in the loop in a 4f configuration. A half wave plate (HWP) was placed in front of the loop optical path to change the power ratio between the beam entering the loop and the beam transmitted straight ahead. The loop output and the straight-through beam were recombined with the PBS, passed through another HWP, and then split by a second PBS into two lateral imaging paths. The reflected s-polarized laser went to Path-2 and the passing p-polarized laser went to Path-1 (**Fig. 1a**). In Path-2, an additional 0.94 m delay optical path was introduced, and two concave lenses were added in this pathway in a 4f configuration. As a result, the four pulses passed through different optical paths with additional delays of 0, 0.94, 1.88, and 2.82 m of additional delay, creating focal points under the objective lens that were delayed by 3.125 ns (**Fig. 1b**).

**Figure 1.**
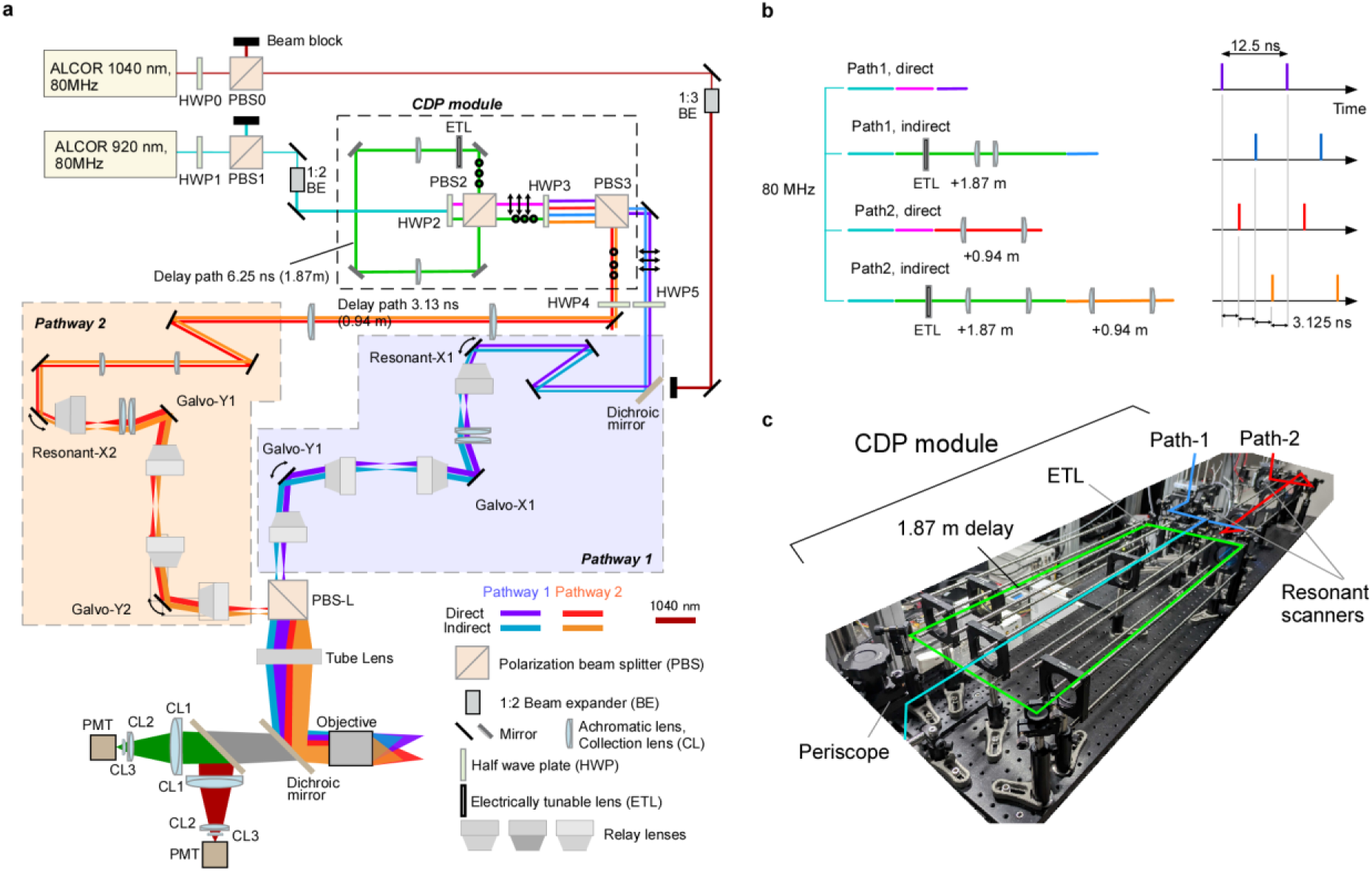
Diesel2p equipped with a CDP module that enables multiplexed imaging along four optical paths. **a.** Schematic diagram of the Diesel2p optical layout with the circular delay-path (CDP) module inserted. **b.** Conceptual drawing of the four paths represented as straight lines, showing spatial scale on the left and temporal scale on the right; each delay is relayed by a dedicated 4 f system, and a 3.125 ns separation between paths 1 and 2 plus a 6.25 ns delay introduced by the CDP module yield an overall separation of 3.125 ns between consecutive paths. **c.** A photograph of the CDP module.

The first and third pulses via Path-1, and the second and fourth pulses via Path-2, scanned separate lateral regions. Within each lateral region, the ETL introduced a focal-depth difference between the two pulses; this depth offset was adjustable. The image sizes of all pathways were the same since ETL was placed at a plane conjugate to the objective pupil. Furthermore, the laser power could be optimized in the two depths and in the two areas by rotating HWPs. This simple loop-based design therefore enabled efficient splitting of an 80 MHz laser into four interleaved beams for multi-depth, multi-area imaging. We call this module, a circular delay-path (**CDP**) module (**Fig. 1c**).

### PSF analysis for four pathways

We first characterized the beam shape in each of the four optical paths by measuring the two-photon point spread function with 0.2 µm beads while the electrically tunable lens was kept at its default setting. For the first direct, first indirect, second direct, and second indirect, we acquired point spread functions (PSFs) at three field positions: the field center, the right edge located 2.5 mm from the center (X-edge), and the upper right corner located 2.8 mm from the center (diagonal-edge). We confirmed that all four paths showed almost identical resolution. Resolution along z-axis was slightly decreased away from the center but the axial full width at half maximum (FWHM) remained below 10 µm. These observations confirm that each field retains the designed optical performance.

### ETL controls focal depth of indirect pathways

Because the first and second indirect paths contain an electrically tunable lens (ETL), their focal depths can be adjusted. We drove each lens from –150 mA to +150 mA and compared the focal depth and PSF full width at half maximum (FWHM). In Path-1I (**Fig. 2c**), the focal depth shifted by ≈0.54 µm per mA. Throughout this range, the lateral FWHM (X and Y) remained virtually unchanged, whereas the axial resolution broadened slightly. Overall, these results show that the ETL provides at least 164 µm of axial translation while preserving effective two-photon excitation efficiency.

**Figure 2.**
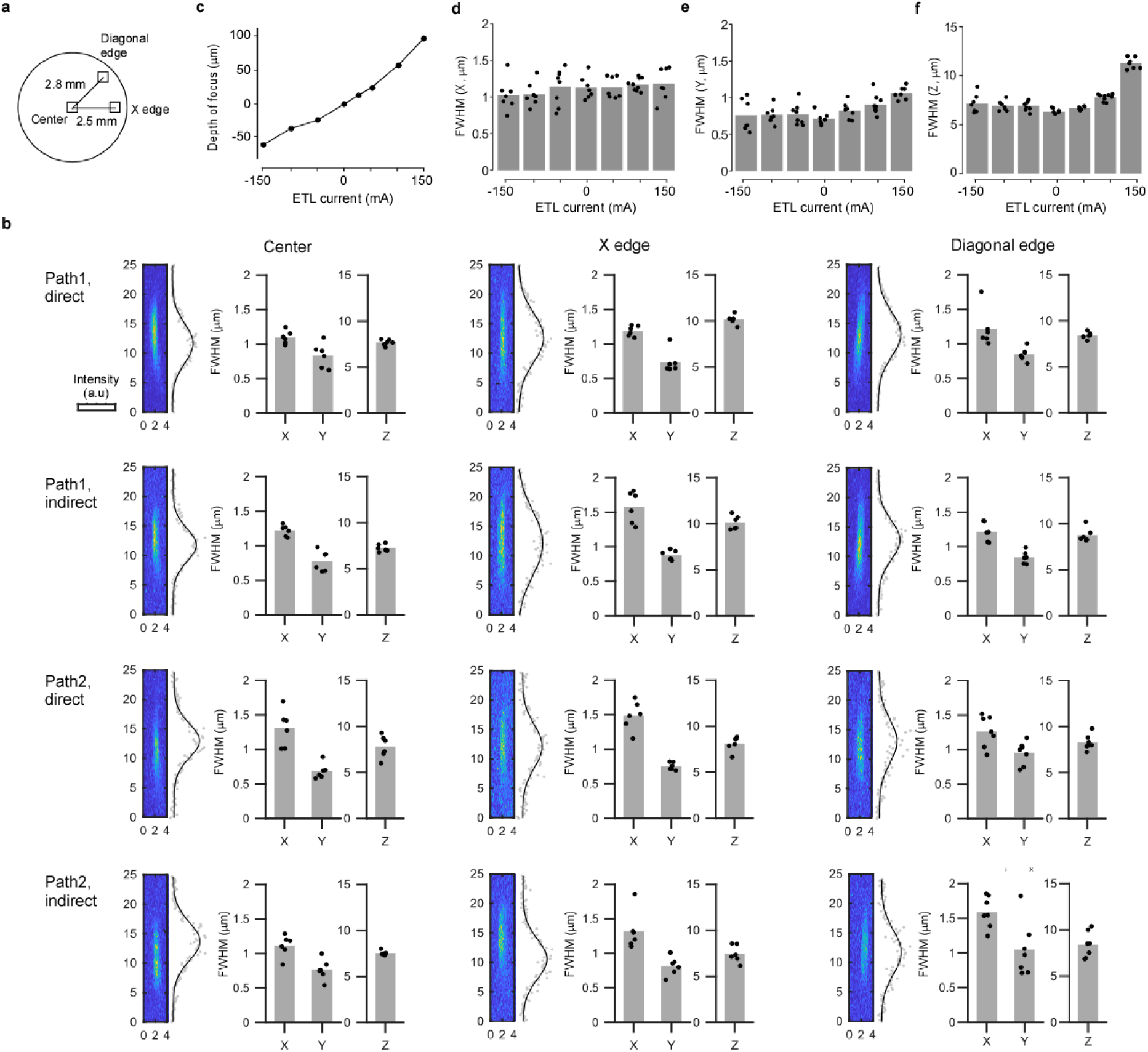
PSF analysis. **a.** Point-spread functions (PSFs) were measured at three locations—image center, X-edge, and diagonal edge—for each of the four optical paths. **b.** The full widths at half maximum (FWHMs) of the PSF in the X, Y, and Z directions are summarized for all four paths using 0.2 µm fluorescent beads; the left sub-panel shows a representative PSF with its axial intensity profile and Gaussian fit, while the right sub-panel lists the X-, Y-, and Z-axis FWHM values (all values in µm). **c.** For path-1D the focal depth was plotted against the driving current of the electrically tunable lens (ETL). **d.** The same data are replotted to show how the horizontal (X,Y) and axial (Z) PSF FWHM varies as a function of ETL current.

### Separating four fluorescence signals using FPGA

To separate the four fluorescence signals on the FPGA we exploited the fixed 3.125 ns delay between successive excitation pulses, which produces an identical 3.125 ns offset in the arrival time of the corresponding fluorescence at the PMT. Before implementing this demultiplexing we verified that individual photon signals were fully contained within that window by imaging the second-harmonic generation of a urea crystal at very low laser power. The PMT output, amplified and filtered identically to in-vivo measurements, was digitized at 3.2 GS/s; most laser pulses produced no response, and a few yielded isolated negative spikes followed by oscillations (**Extended Data Fig. 1a–c**). The negative peak had a full width at half maximum of 1.8 ns (**Extended Data Fig. 1d**), safely shorter than 3.125 ns, and the timing of the first trough and second positive peak was highly correlated across pulses (r = 0.77; **Extended Data Fig. 1e**), indicating that the oscillation originated in the detection electronics rather than in photon statistics. Because this oscillatory tail was much smaller than the initial spike it could be rejected by integrating within a suitably short window. Accordingly, the FPGA assigned four non-overlapping 3.125 ns integration windows to the photomultiplier output, summed the photon counts for each pulse in real time, and streamed the results as four independent channels corresponding to Path-1D, Path-1I, Path-2D, and Path-2I. This pulse-by-pulse demultiplexing removed temporal cross-talk before image reconstruction and allowed simultaneous acquisition of four focal planes without sacrificing temporal resolution.

**Extended Data Fig. 1.**
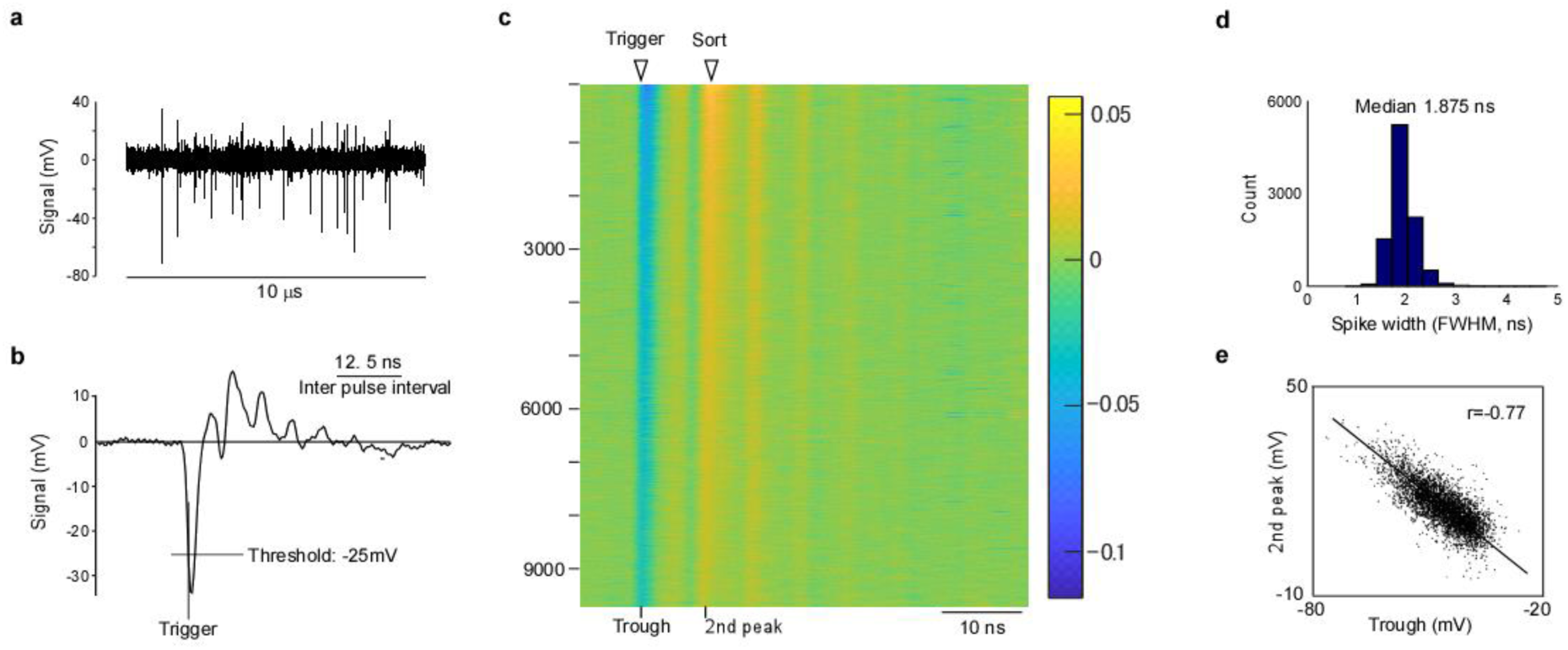
Single photon signal. **a.** Analog PMT signal sampled at 3.2 GS/s over 10 µs; the urea crystal was excited at the minimum power that still produced detectable fluorescence, so each negative pulse is taken to represent a single photon. **b.** Average waveform of the negative pulses in a, showing a damped oscillation that follows the initial trough. **c.** Color map of ≈10 000 aligned pulses used for the average in b, sorted by the amplitude of the second positive peak; nearly every pulse exhibits the same temporal oscillation. **d.** Distribution of the full width at half maximum of the negative pulses; the median is 1.875 ns. **e.** Scatter plot of the time bins containing the first trough and the second positive peak for each pulse, revealing a strong correlation (r = 0.77).

### FPGA system

The purpose of our system is to distinguish in the time domain the four fluorescence signals that follow four excitation pulses spaced 3.125 ns apart (**Fig. 3a**). Some experiments do not need all four optical paths and often do not require single-pulse resolution, so the module can operate in several modes that compress the data stream and lighten later analysis (**Fig. 3b**). Users set three parameters: the number of active channels, which temporal bins each path should sum, and the accumulation count that determines how many consecutive pulses are averaged for compression.

**Figure 3.**
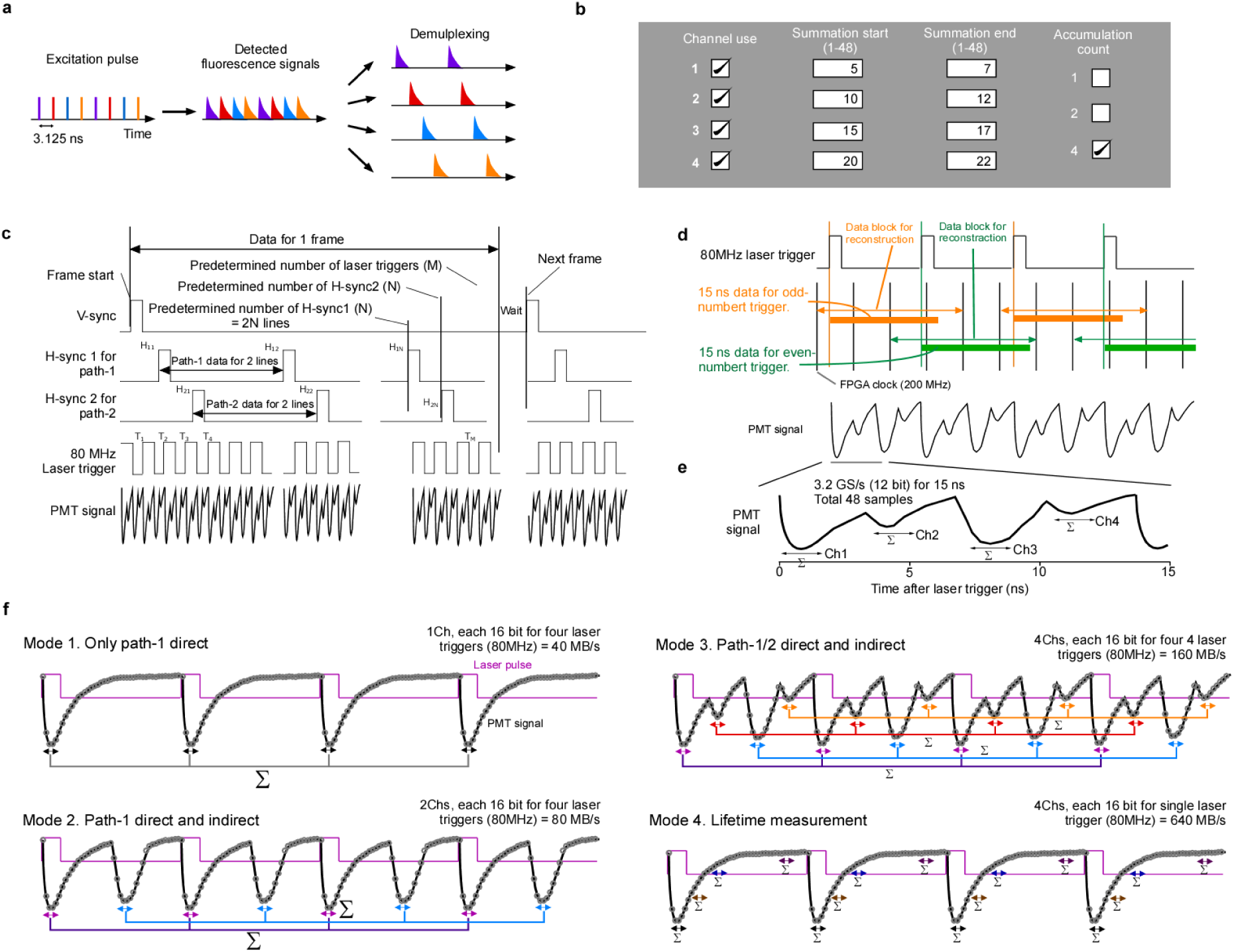
Operating modes of the FPGA digitizer and the underlying mechanism. **a.** The system separates four fluorescence signals that arrive 3.125 ns apart. **b.** The FPGA digitizer operates in several configurations; here it runs in the mode that uses all four optical paths and sums data from four successive pulses. **c.** A frame is defined by five inputs—vertical sync, two horizontal sync signals, the laser clock, and the PMT output. After a vertical-sync pulse marks frame start, the module acquires PMT data for the prescribed numbers of horizontal-sync events (N) and laser pulses (M). **d.** Signal processing for odd and even laser pulses is interleaved (orange for odd, green for even). Forty-eight samples are stored per pulse, covering 15 ns so that fluorescence lifetimes extending beyond the 12.5 ns pulse period are fully captured. **e.** Representative PMT waveforms recorded for the four optical paths. **f.** Operating modes: Mode 1 records a single path and, by summing four pulses, yields 20 MS/s; Mode 2 records the direct and delayed branches of path 1; Mode 3 records all four paths simultaneously; Mode 4 records four-gate lifetime data.

To verify single-photon signals we first recorded at 3.2 GS/s, but at that rate the two-gigabyte on-board memory fills almost immediately. To allow continuous recording the FPGA integrates the photomultiplier output within four windows for every eighty-megahertz pulse and stores four sixteen-bit values (**Fig. 3c**). The module receives five inputs simultaneously: the laser-clock TTL and the photomultiplier waveform at 3.2 GS/s, two horizontal sync signals from the resonant scanners at 200 MHz, and a vertical sync signal at 200 MHz that marks each frame start. It outputs only the timing of the horizontal syncs relative to the vertical sync and the four photomultiplier values recorded at fixed delays after each pulse. With these data four images can be reconstructed. Scanner control and image formation remain in separate software, which makes integration with any commercial microscope straightforward.

Full details of the subVIs appear in the **Supplementary Information**. Here we describe how partial duplication and compression prevent data loss. Because some fluorescence arriving eleven nanoseconds after a pulse must be integrated up to fourteen nanoseconds when the lifetime is three nanoseconds, information from the next pulse is needed. We duplicated the arithmetic cores so that odd and even pulses are processed in parallel (**Fig. 3d,e**). Each pulse is handled over fifteen nanoseconds, forty-eight samples in total, without interruption.

After integration the FPGA outputs four sixteen-bit numbers for every eighty-megahertz pulse. This creates a data stream of 640 MB/s. An optional accumulation count allows up to four successive pulses to be summed before transfer (**Fi3. 4b**). The per-channel rate then becomes 20 MHz, comparable to a conventional two-photon microscope. Diesel2p contains two autonomous resonant scanners that cannot be hardware-synchronized, so separate software associates the condensed photomultiplier data with the two scanner triggers in real time. Any user can add the FPGA module and its host PC to an existing microscope and obtain multiplexed acquisition. Five representative operating modes are shown in **Fig. 3f**.

### Modular hardware and software

To keep the system compatible with many microscopes we adopted a modular design. The hardware accepts only synchronization and photomultiplier signals. The software performs real-time accumulation on the FPGA, sends the reduced data via Thunderbolt, and saves it to disk. Scanner control stays in the existing microscope software, so we could switch between the original controller and the high-speed sampler simply by reconnecting BNC cables. Each subVI carries out a single task, so pulse rate, acquisition speed, or spectral multiplexing can be adjusted by replacing one module. Adding this unit to a running two-photon microscope immediately enables laser multiplexing and lifetime imaging.

### Temporal separation of GCaMP signals from distinct pathways

To image GCaMP fluorescence on four channels simultaneously, we first quantified inter-channel leakage using acute cortical slices (**Fig. 4a,b**). With only the Path-1D channel active, the mean GCaMP waveform clearly spilled into the other channels. In the in vivo configuration, the four pathways are sampled 3.125 ns apart (10 digitizer bins). Therefore, when GCaMP peaks on Path-1D, its leakage appears sequentially on Path-2D, Path-1I, and Path-2I at +3.125, +6.250, and +9.375 ns, respectively; by symmetry, the same ordering holds for any source channel (e.g., Path-2D leaks to Path-1I, Path-2I, then Path-1D). Let the true images be *I* = [*I*1, *I*2, *I*3, *I*4]⊤ and the measured images *J* = [*J*1, *J*2, *J*3, *J*4]⊤. Then

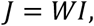

**Figure 4.**
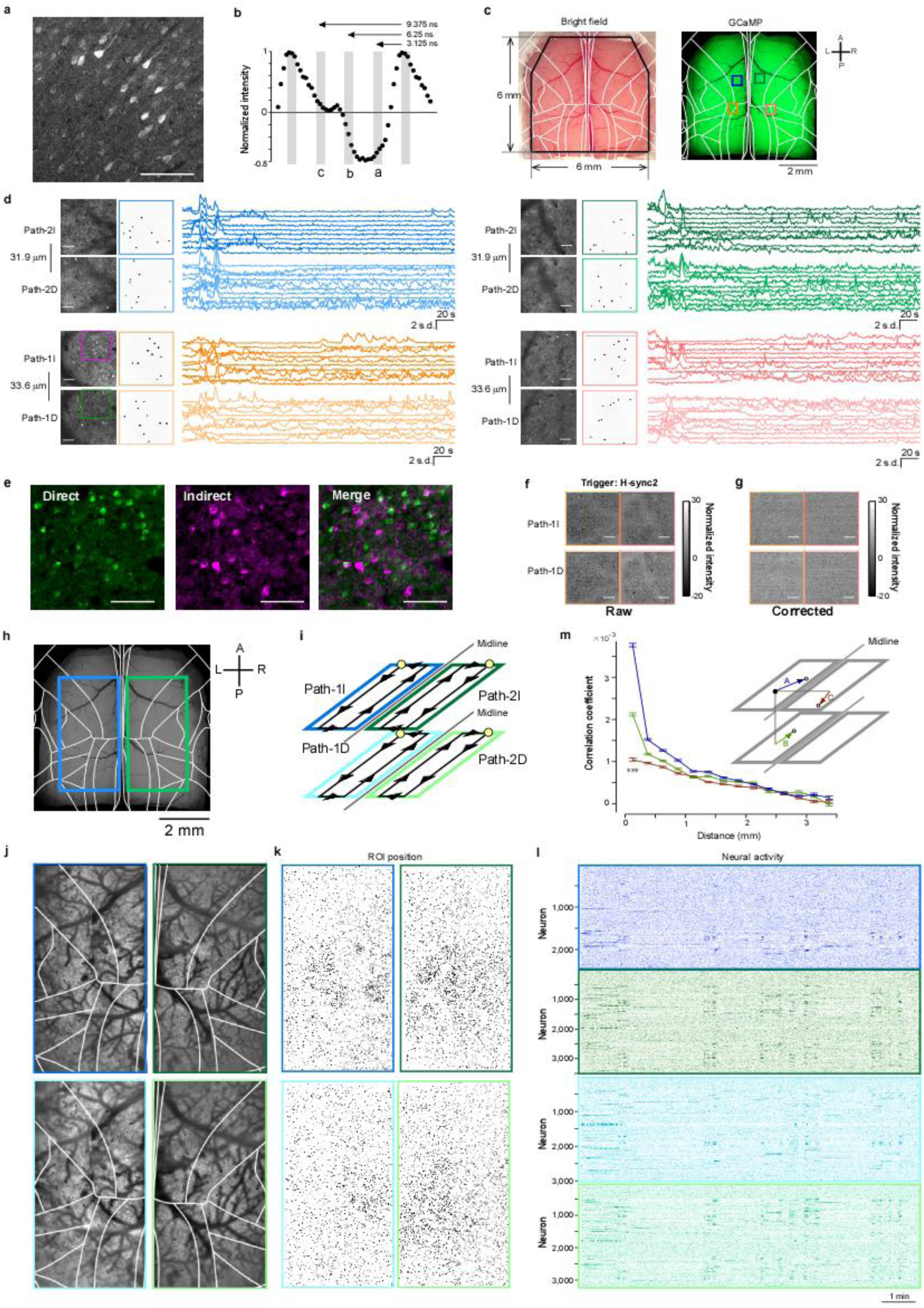
Four pathways two-photon calcium imaging. **a.** Two-photon image of a cortical slice containing GCaMP-expressing pyramidal neurons. Scale bar, 100 µm. **b.** Normalized fluorescence as a function of time bins within the 12.5-ns laser period. Baseline was taken from a background region without tissue and traces were scaled by the maximum amplitude. Three non-overlapping windows (a–c), spaced by 3.125 ns, were used to estimate and correct inter-channel leakage. **c.** Dorsal view of mouse cortex with a 6 mm × 6 mm cranial window (left) and GCaMP epifluorescence (right). Colored squares mark the four fields used for simultaneous four-area/two-depth imaging shown in **d**. **d.** Example simultaneous calcium imaging from four areas at two depths. Signals were corrected for inter-channel leakage using coefficients derived from **b**. Ten ROIs were randomly chosen per field and their fluorescence time courses are shown; colors match **c**. Each field of view (FOV) was 0.5 mm × 0.5 mm, acquired at 15 Hz for 5 min. The direct pathways imaged the deeper plane of each pair. Laser power: Path-1D, 115mW, Path-1I, 105mW, Path-2D, 122mW, Path-2I, 110mW. Imaged at 15 frame/s. **e.** Direct (green) and Indirect (magenta) channels, separated by Δt = 6.25 ns, show negligible cross-talk (merge at right). **f.** To assess leakage between adjacent bins (Δt = 3.125 ns), Path-1 raw signals were reconstructed using the H-sync-2 trigger; mean of 100 frames shown. Bright vessels and dark somata reflect sign-reversed leakage. **g.** Same analysis after leakage correction; vessel/soma leakage is no longer visible. **h.** Bilateral two-layer imaging patches overlaid on atlas. Scale bar, 2 mm. **i.** Schematic of the four pathways (Path-1D, Path-1I, Path-2D, Path-2I). Yellow circles indicate scan start points; black lines/arrows indicate the galvanometric slow-scan trajectory. The resonant scanner sweeps orthogonal to these lines (not drawn). Each pathway images a distinct layer in one hemisphere. **j.** Mean two-photon images for the two depths in both hemispheres (top and bottom panels are different depths of the same areas). Box colors correspond to pathways. Laser power: Path-1D, 115mW, Path-1I, 105mW, Path-2D, 122mW, Path-2I, 110mW. Imaged at 3.6 frame/s. **k.** ROIs detected in **j**. **l.** Fluorescence time courses from ROIs in **k**, ordered by anterior-to-posterior position within each FOV. **m.** Pairwise correlation coefficients plotted against lateral distance between ROIs. Blue, pairs within the same FOV; green, pairs across the two depths in the same hemisphere (z-separation of 32 µm ignored so distance is lateral only); red, inter-hemispheric pairs computed after reflecting one ROI across the midline. Error bars denote s.e.m. The closest distance bin (<0.25 mm) showed significant differences among groups (Wilcoxon rank-sum test, *p* < 0.001).

where W is a circulant matrix determined by the leakage coefficients *a*, *b*, *c* defined in Fig. 5b. The demixed estimate of the true signals is obtained by

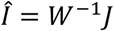

**Figure 5.**
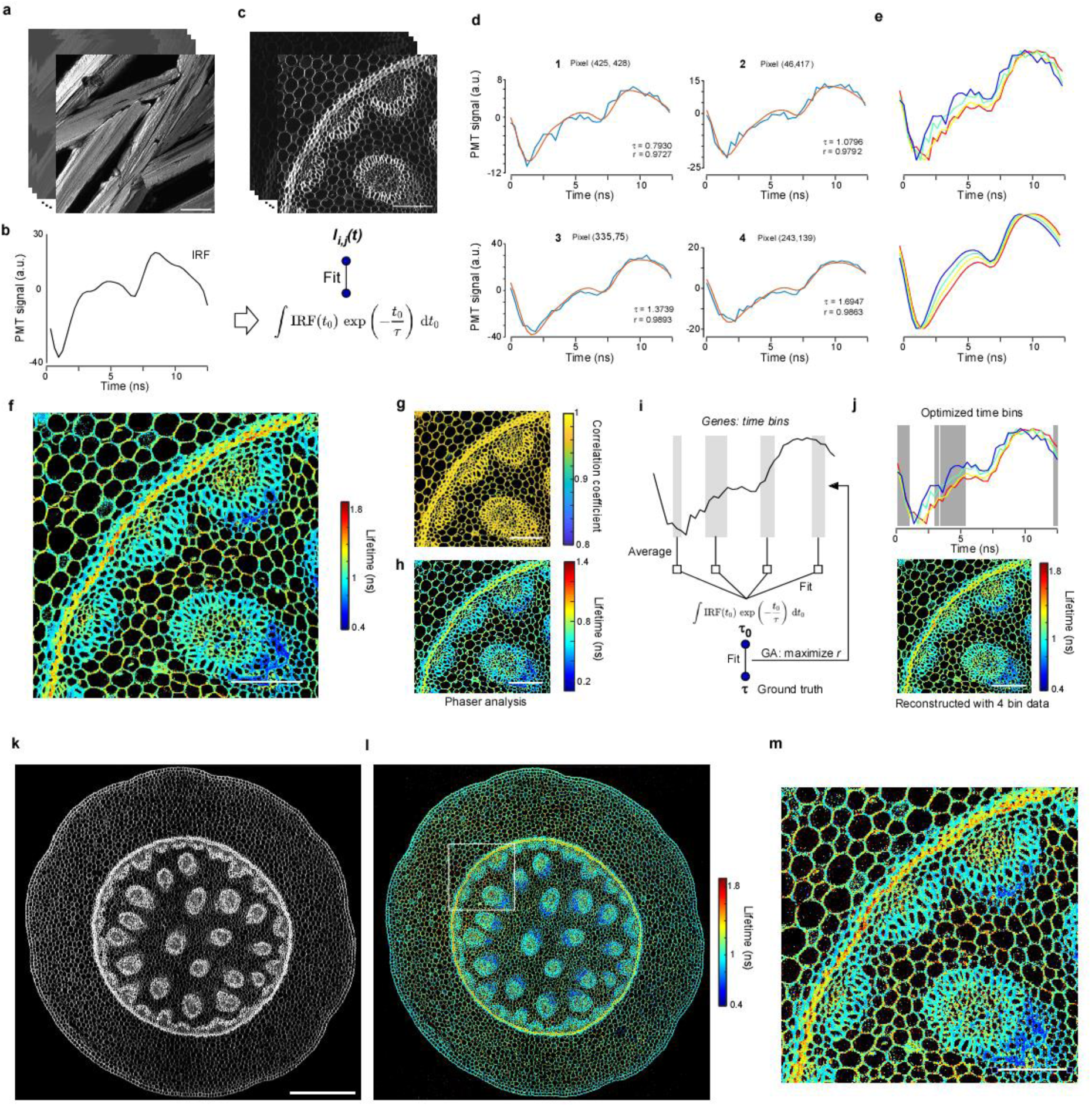
Large FOV two-photon lifetime imaging with Diesel2p. **a.** Two-photon imaging of urea crystals. The signal is second-harmonic generation and therefore represents the system impulse response function (IRF). Forty consecutive images visualize the 12.5 ns pulse period. Scale bar, 100 µm. **b.** Average of the forty images in **a**, giving the IRF. **c.** Two-photon fluorescence imaging of Convallaria, top panel. Forty consecutive images visualize the 12.5 ns pulse period. In the bottom panel each pixel intensity trace is fitted with the convolution of the IRF in b and a single-exponential decay, yielding the fluorescence lifetime *τ*. **d.** Temporal traces and fits for four representative pixels taken from **c**. **e.** Overlay of the four traces and fits shown in **d**, colored according to the lifetime scale in **f**. **f.** Lifetime map in which pixels brighter than background are colored by their fitted *τ* value. Scale bar, 100 µm. **g.** Pixel-wise correlation coefficient of each fit in **f**. Most pixels show a value greater than 0.95. Scale bar, 100 µm. **h.** Lifetime image obtained by phasor analysis of the same data set. Scale bar, 100 µm. **i.** Procedure for selecting four time-bins that allow lifetime estimation from only four measurements. A genetic algorithm optimized the bin positions using pixels in the lower-left triangle of the image. **j.** Lifetime image produced with the four optimized bins from **i**. High accuracy was obtained even in the upper-right triangle, which the algorithm did not use for training. Scale bar, 100 µm. **k.** Large FOV two-photon image of the entire Convallaria section. Scale bar, 500 µm. **l.** Lifetime map corresponding to **k**. **m.** Magnified view of the boxed region in **l**; this region corresponds to panel **f, h,** and **j**. High estimation accuracy was retained across the large field of view. Scale bar, 100 µm.

(see **Methods** for detail).

### Simultaneous four-area, two-plane calcium imaging in vivo

To test the demultiplexing in vivo, we used double-transgenic mice expressing GCaMP6s broadly in cortical excitatory neurons (Ai162 × Slc17a7-IRES2-Cre). Mice were head-fixed under a 6 mm × 6 mm cranial window; dorsal epifluorescence confirmed widespread expression (**Fig. 4c**). We targeted four 0.5 mm × 0.5 mm fields across the two hemispheres. Each pathway of the path1/2 imaged two of the fields, with the direct plane positioned deeper than the indirect plane by ≈32 µm. Thus, eight images (four areas × two depths) were acquired simultaneously at 15 Hz for 5 min (**Fig. 4d**). Ten neurons per field were randomly selected to display fluorescence time courses. Visual inspection of magnified mean images revealed negligible cross-talk between the Direct and Indirect channels separated by 6.25 ns (**Fig. 4e**). Because adjacent channels are offset by only 3.125 ns, we also tested leakage between neighboring bins by reconstructing Path-1 frames using the Path-2 trigger (H-sync-2). This produced sign-reversed vessel and soma contrast on Path-1, indicating opposite-phase leakage (**Fig. 4f**). After applying the leakage correction (*W*^−1^), the sign-reversed image disappeared (**Fig. 4g**), confirming accurate removal of cross-talk between adjacent bins.

### Bilateral large field-of-view two-layer imaging

We next expanded the field of view to cover large portions of dorsal cortex bilaterally in awake mice (**Fig. 4h**). During each slow-scan cycle of the galvanometers, each pathway scanned one of two depths in one hemisphere (**Fig. 4i**). Representative mean two-photon images (**Fig. 4j**), corresponding ROIs (**Fig. 4k**), and their fluorescence time series (**Fig. 4l**) are shown. Pairwise correlations of neural activity decreased with lateral distance (**Fig. 4m**). Within-FOV, same-layer pairs showed the highest correlations, followed by within-FOV, different-layer pairs (z-offset ≈ 32 µm, distance computed laterally), and then interhemispheric pairs computed after reflecting one ROI across the midline. The closest distance bin (< 0.25 mm) showed significant differences between groups (Wilcoxon rank-sum test, p<0.001). These results are consistent with columnar organization and interhemispheric symmetry.

### Bilateral four-layer imaging

Finally, to image four distinct depths simultaneously, we introduced a focal offset between Path-1 and Path-2 by translating the paired achromatic doublet placed immediately upstream of the Path-2 X-galvo. Combined with depth tuning of the Indirect paths via the electrically tunable lens, this yielded four independent focal planes (**Extended Data Fig. 2a,b**). Using this configuration, we recorded four layers across medial regions of both hemispheres simultaneously (**Extended Data Fig. 2c–e**).

**Extended Data Fig. 2.**
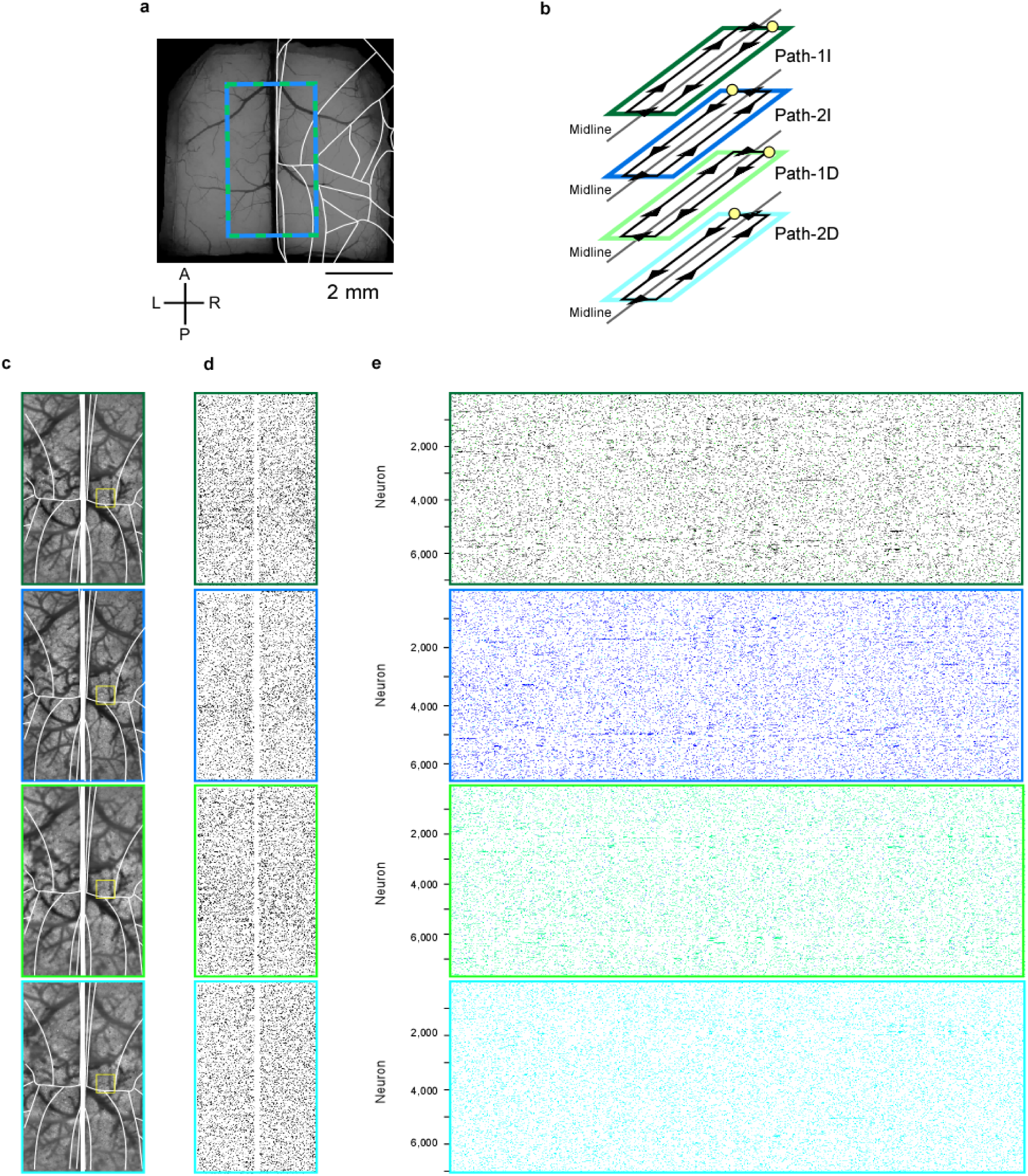
Bilateral large field-of-view four-layer two-photon imaging. **a.** Imaging footprint overlaid on dorsal vasculature and the Allen Mouse Brain Atlas. Axes as indicated. Scale bar, 2 mm. **b.** Acquisition schematic. The four pathways (Path-1D, Path-1I, Path-2D, Path-2I) each scan a distinct cortical layer across hemispheres. Yellow circles mark scan start-points; black lines/arrows indicate the galvanometric slow-scan trajectory. The resonant scanner moves orthogonally (not drawn). Together, the pathways provide bilateral coverage at two depths per hemisphere (four layers total). **c.** Mean two-photon images from the same lateral position at four depths (corresponding to **d,e**). Frame colors map to the pathway identity. Laser power: Path-1D, 115mW, Path-1I, 105mW, Path-2D, 122mW, Path-2I, 110mW. Imaged at 3.6 frame/s. **d.** ROIs detected from the fields in **c**. **e.** Fluorescence time courses for the ROIs in **d** (one row per ROI), ordered by the anterior-to-posterior position within each field.

### Large FOV 2pFLIM

We performed two-photon fluorescence lifetime imaging (2p-FLIM) using a 3.2 GS/s high-speed digitizer (**Fig. 3f**). Sections of Convallaria, whose endogenous fluorescence lifetime mirrors tissue architecture, are widely used as FLIM standards. In our setup, the 12.5 ns interval between successive 80 MHz laser pulses was divided into 40 temporal bins. Consequently, we first acquired a stack of 40 images with the same FOV, each averaged over 1,000 frames. Comparing pixel intensities across these 40 frames yielded fluorescence signals with a temporal resolution of 12.5 ns / 40. Because the system can store data from only four bins per acquisition, imaging had to be repeated ten times to obtain the full 40-bin dataset. We then characterized the system response using the second-harmonic generation (SHG) signal from urea crystals (**Fig. 5a,b**). As SHG is generated with zero temporal lag, it provides the instrument response function (IRF). Convolving this IRF with a single-exponential decay of lifetime τ and fitting it to the Convallaria data allowed us to determine the fluorescence lifetime of each pixel (**Fig. 5c–e**). The Convallaria signals were accurately described by the IRF-convolution model, enabling resolution of subtle lifetime differences. Applying this fitting to all pixels produced a fluorescence-lifetime image (**Fig. 5f**); most pixels exhibited a correlation coefficient > 0.95 between the model and the data (**Fig. 5g**). Phasor-plot analysis yielded an equivalent lifetime map, aside from a slight phase shift (**Fig. 5h**). Together, these results confirm that our FPGA-based system is well suited for lifetime imaging.

To obtain a lifetime image in a single scan, we optimized the choice of four time-bins available per excitation pulse. A genetic algorithm was used to identify the four-bin combination that best reproduced the lifetimes determined from the complete 40-bin dataset (**Fig. 5i**). The selected bins generated lifetime images virtually indistinguishable from the 40-bin reference (**Fig. 5j**).

Finally, the field of view was enlarged to encompass the entire Convallaria section, and images of comparable quality were obtained (**Fig. 5k–m**). These findings demonstrate that integrating fluorescence lifetime imaging with the Diesel2p mesoscale microscope enables simultaneous acquisition of wide-field lifetime data.

## Discussion

In this study, we split the 80 MHz laser beam into four paths, enabling four-field mesoscale imaging with the widely used 80 MHz light source. We achieved efficient, loss-free signal processing by subdividing the 12.5 ns inter-pulse interval into 40 bins (0.31 ns each) and compressing the data on the fly with an FPGA. Using this system, we were able to perform large-scale calcium imaging in vivo of the bilateral wide dorsal cortex across up to four planes. We also demonstrated lifetime imaging, confirming sub-nanosecond resolution using Convallaria sections with a large field of view (FOV).

Although the Diesel2p is the only mesoscale microscope with an open-source optical path, we have additionally released fully open-source control software and acquisition hardware. Our new FPGA system has a modular architecture that is independent of the scanner-control electronics, so it can be plugged into any existing two-photon microscope via a BNC cable and used immediately. Multiplexing and lifetime imaging are therefore available not only on Diesel2p but also on conventional systems, and compatibility with the ubiquitous 80 MHz laser lowers the adoption barrier. Excitation efficiency on Diesel2p is close to its power limit with four fields, but further multiplexing should be possible with a lower-repetition-rate laser or adaptive optics. While we do not achieve the temporal resolution of TCSPC, the ability to record multiple FOVs or large FOV simultaneously offers a significant increase in acquisition speed. The FPGA code is clearly modular and written in the standard National Instruments LabVIEW software, which facilitates extensions such as machine-learning-based image analysis (Song et al. 2023; Stringer et al. 2020), integration with optical stimulation, and closed-loop or BMI experiments (Hira et al. 2014; Hira 2024).

The system developed by Jerry Chen’s group (Clough et al. 2021) used two laser colors; our design also supports dual-color excitation, making palette expansion straightforward. Techniques such as reverberation and light-beads can deliver even higher multiplexing, but when Diesel2p is driven by an 80 MHz laser, four beams are the practical limit, making our system a realistic best solution. Combining Diesel2p with a reverberation circuit, a light-beads module (Demas et al. 2021; Beaulieu et al. 2020), or a FACED module (Lai et al. 2021) is nevertheless possible. For instance, lowering the repetition rate to ≈10 MHz would theoretically allow an eight-fold increase, yielding 32 beams—likely close to the in vivo limit for two-photon excitation. Multi-pixel detectors such as SiPM arrays can mitigate crosstalk and raise the ceiling further, yet laser-induced brain damage will ultimately constrain in vivo power.

With FLIM now feasible on a large FOV two-photon microscope, a diverse set of FRET sensors including cAMP (Massengill et al. 2022), PKA (Ma et al. 2022), CREB (Laviv et al. 2020), CaMKII (Jain et al. 2024), Ras-GTPase (Yasuda et al. 2006; Murakoshi, Wang, and Yasuda 2011), ERK/MAPK (Hirata and Kiyokawa 2019), and PI3K/PTEN (Kagan et al. 2025) can be applied to large-scale in vivo brain imaging. Because FLIM is robust against biological motion, it is valuable not only for calcium indicators with pronounced lifetime changes (van der Linden et al. 2021) but potentially also for membrane-voltage sensors (Bowman et al. 2023). One can conduct macroscale RCaMP imaging on optical Path-1 while simultaneously performing spine-level cAMP or Ras FLIM imaging on Path-2, bridging macroscale activity with local biochemistry across multiple time scales. The combination of large FOV coverage and lifetime imaging should accelerate multi-signal detection in cleared brains, speeding transcriptomic and connectomic mapping exponentially. Label-free measurements of NAD(P)H and FAD (Georgakoudi and Quinn 2023; Yaseen et al. 2013) and separation of SHG signals (Zipfel et al. 2003) are also possible. Beyond the brain, in vivo FLIM has been demonstrated in retina (Walters et al. 2022), lung (Ryu et al. 2018), intestine (Quansah et al. 2023), pancreas (Azzarello et al. 2022), liver and kidney (Lin et al. 2022), with rapid progress toward human applications. In disease research, the platform enables studies of cancer (Ouyang et al. 2021), Alzheimer’s disease (Norambuena et al. 2024), diabetes (Wang et al. 2023), the immune system (Paillon et al. 2024), and vascular disorders. Thus, broad adoption of our open-source system promises wide-ranging contributions from basic science to clinical medicine.

## Acknowledgements

We thank H. Kasai, K. Ichige, Y. Kimura, S. Kimura, H. Kurosu, R. Hattori, S. Tanno, O. Ooishi for technical advice and discussion. This work was supported by JP22wm0525007 (RH), JP25wm0625405 (RH), JP19dm0207089 (YI), JP23wm0525008 (AF) from AMED, JP22H02731 (RH), JP20K22678 (RH), JP21B304 (RH), JP21H05134 (RH), JP21H05135 (RH), JP21H05242 (YI), JP23H02589 (YI), JP25H02599 (AF), JP24H02150 (AF) JP21H05243 (AF), JP24KK0186 (AF, RH) from MEXT/JSPS, JPMJFR231X (RH), JPMJCR1751 (YI) from JST, Nakatani Foundation (RH), Shimadzu Foundation (RH), Takeda Science Foundation (RH), The Precise Measurement Technology Promotion Foundation (RH), The Sumitomo Foundation (AF, RH),Tateishi Science and Technology Foundation (RH), and Research Foundation for OptoScience and Technology (RH).

## Author contributions

Riichiro H. conceived the project. Riichiro H. H.I. F.I. Y.Y. conducted optics experiments. H.I. F.I. optimized the large cranial window surgery. H.I. F.I. Y.Y. T.H conducted imaging experiments. Reiko H. optimized imaging environments. Riichiro H. O.F. A.K. A.F. built FPGA-based imaging software. Riichiro H. S.S. designed a new version of Diesel2p. C-H.Y. S.L.S. supported the optics. Riichiro H. H.I. F.I. T.H analyzed the data. S.K. K.K. produced the AAV. Y.I. H.S. H.T. A.F. supervised the project. Riichiro H. wrote the paper with the comments of all authors.

---

**Supplementary Fig. 4.**
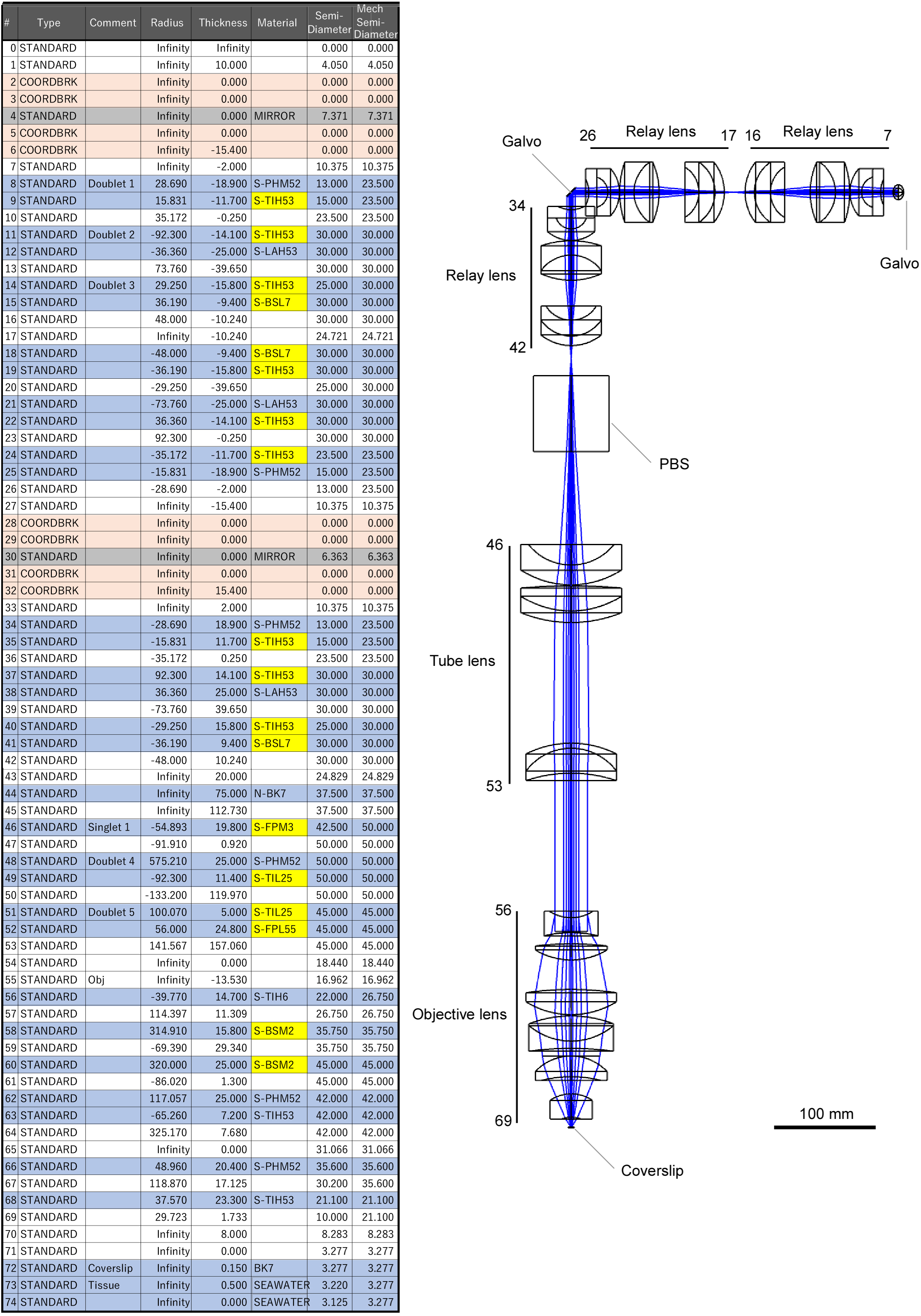

## Notes

### Competing Interest Statement

The authors have declared no competing interest.

