## Supplementary information for "Open-source modular FPGA system for two-photon mesoscope enabling multi-layer, multi-depth neural activity recording and lifetime imaging"

The supplementary information includes 1. Supplementary discussion, 2. FPGA operation mechanisms, 3. Instruction of the LabVIEW software, and 4. Modified optical parameters for Diesel2p.

### 1. Supplementary discussion

Two strategies have been used in laser multiplexing: signal separation using analog electronics and high-speed digital processing. For example, in Trepan2p (Stirman et al. 2016; Yu et al. 2022), to bifurcate the PMT signal, the signal was converted to two TTL signal trains by the ultra-fast discriminator (TD3000) during the two 6.25 ns time windows. The time windows were generated by the other two discriminators after inserting a delay box between the 80 MHz sync signal of the laser. The conversion of the PMT signal to TTL signal train using a fast discriminator is equivalent to counting single photons. There is little loss in case of the dark sample. The two TTL signals were then recorded at about 10 M samples per second after low-pass filtering of the TTL signals with ≈10MHz cut-off frequency. In the paper by Cheng and colleagues (Cheng et al. 2011), a circuit was designed to amplify signals in four time windows using a 3-ns pulse synchronized to the laser output and an analog multiplier (ADL5391, Analog Devices) (Supplementary Figure 4 of (Cheng et al. 2011)). While these designs for high-speed sampling can aim for the highest performance in individual systems, they require in-depth knowledge of optics in analog electronic circuitries.

High-speed digital processing can minimize knowledge of analog electronic circuits. For example, in the system developed by Clough and colleagues for Diesel2p, the PMT signal was digitized with a NI-6587 (1 GS/s) and the signal was processed at high-speed with a PXIe-7691R (FPGA) (Clough et al. 2021). This strategy is equivalent to digitizing the process using a high-speed discriminator circuit as used in original Diesel2p. Beaulieu and colleagues combined the NI5771 with a PXIe-7-1972 (2G onboard memory) FPGA, which can analog record PMT signals at 3.0 GS/s (8bit) (Beaulieu et al. 2020). They used a pulse picker to reduce the laser repetition rate to 20 MHz, and an Analog Devices AD9516 to generate the digitizer clock. Demas and colleagues are similar to Beaullier and colleagues (Demas et al. 2021). They recorded at 1.6GS/s, 12bit with NI 5771 and FPGA PXIe-7575R (2G onboard memory). Wu et al. did almost the same, using PXIe-5160 (2G onboard memory, National Instruments) and getting 625 MS/s (Wu et al. 2020). These three labs also customized Vidrio Inc's ScanImage software platform, although this software is not open source. It also differs from our method in that it uses a lower laser repetition frequency and hardware to generate the digitizer clock from the timing of the laser emission. Our method does not assume synchronization of the laser output signal with the FPGA clock. This allows us to simply add the FPGA circuitry module to the conventional two-photon microscopy.

### 2. FPGA operation mechanisms

#### 2-1. Signal processing performed for each V-Sync detection

We referred to the frame start signal as the vertical synchronization (V-sync) signal, and the synchronization signal from each resonant scanner as the horizontal synchronization (H-sync) signal. The Diesel2p system is equipped with two resonant scanners for Path-1 and Path-2; therefore, H-sync1 and H-sync2 were recorded separately for each pathway.

The V-sync signal was detected at a sampling rate of 200 MS/s. Upon detection of the rising edge of the V-sync signal, we began counting the number of H-sync1 and H-sync2 pulses and recorded the elapsed time from the onset of V-sync at each detection. The total number of H-sync1 and H-sync2 pulses to be detected was pre-defined and corresponded to half the number of vertical scan lines per frame (i.e., the number of round trips of the resonant scanner).

Figure 4 illustrates the processing loop: after acquiring H-sync1 and H-sync2 signals for N repetitions, the system entered a wait state until the next V-sync edge, at which point the counting resumed.

In parallel, following the detection of each V-sync edge, PMT signal acquisition and processing were performed for each 80 MHz laser pulse over a finite time window. This procedure was repeated M times, where M defines the number of laser pulse–associated signals collected within a single frame. For instance, if the number of image lines generated by pathway 1 is 1600, then N equals 800; since an 8 kHz resonant scanner completes 800 round trips in 0.1 seconds, this corresponds to a 0.1-second acquisition period. During this time, an 80 MHz laser emits eight million pulses, setting the maximum value of M to 8,000,000. The horizontal width of the resulting image is determined by the number of PMT signals averaged over the corresponding number of laser pulses, as described later.

#### 2-2. Measurements for sync signals

The Diesel2p system employs two resonant scanners for pathway 1 and pathway 2, necessitating independent recording of H-sync1 and H-sync2. Upon detection of a V-sync signal, counting of H-sync1 and H-sync2 was initiated. H-sync1 and H-sync2 signals were sampled at 200 MS/s. Each time an H-sync1 or H-sync2 pulse was detected, the total trigger count was incremented, and the elapsed time since the corresponding V-sync signal was buffered. Separate buffers were used to independently process H-sync1 and H-sync2 signals. The buffered trigger counts and corresponding elapsed times were then sequentially transferred to the PC.

Counting stopped once a predefined number of H-sync1 and H-sync2 pulses had been recorded, after which the system awaited the next V-sync edge. Compared to the data volume from the PMT, the transfer bandwidth required for these sync signals was relatively small. The H-sync frequency was approximately 8 kHz, resulting in 8,000 samples of elapsed time (32-bit) and trigger count (32-bit) per V-sync interval. If a V-sync signal arrived while a finite number of H-sync1 or H-sync2 pulses were still being processed, that V-sync signal was skipped. The frequencies of H-sync1 and H-sync2 were nearly identical, as both were driven by resonant scanners with the same frequencies.

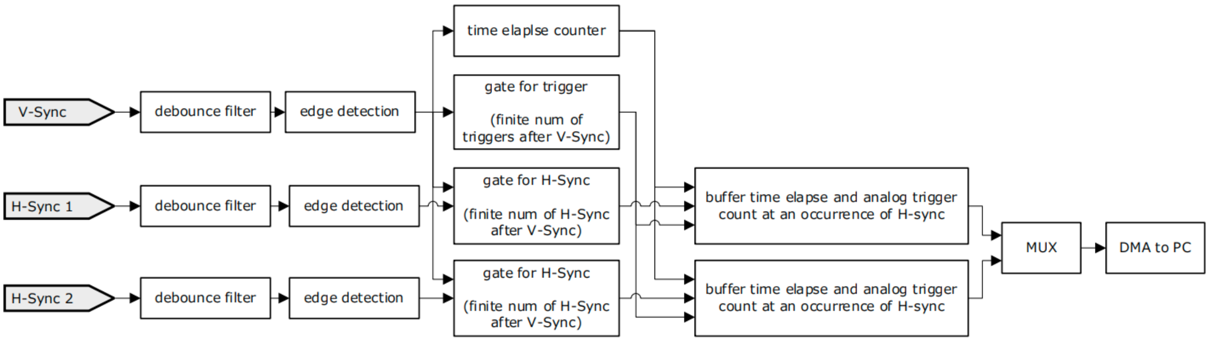


**Supplementary figure 1 | Architecture of the trigger-counting and time-stamping circuit driven by V-Sync / H-Sync signals**

V-Sync and two H-Sync signals first pass through debounce filters to remove chatter, then edge detectors register their rising edges. A time-elapse counter, reset on the rising edge of V-Sync and shared across all paths, serves as the time reference. After each V-Sync, a trigger gate that is active for only a limited number of events opens, while each H-Sync path has an H-Sync gate that opens for a predefined number of H-Sync pulses. Whenever an H-Sync pulse is detected within its valid window, the current counter value and the number of analog triggers observed during that H-Sync period are stored as a pair in a buffer. Outputs from multiple buffers are combined by a multiplexer (MUX) and transferred to the PC via high-speed direct memory access (DMA), enabling real-time acquisition of precisely synchronized trigger timestamps and counts.

**2-3. Parallelized process to acquire 48 samples at every 80MHz trigger**

The 80 MHz laser trigger signal arrived every 12.5 ns. The FPGA operated at 3.2 GS/s, corresponding to 40 samples per 12.5 ns. Data acquisition was performed in blocks of 16 samples, resulting in an effective FPGA clock frequency of 200 MHz. It should be noted that the 80 MHz laser trigger signal and the 200 MHz FPGA clock were not synchronized. As a result, the timing of the laser trigger varied relative to the start phase of the FPGA clock. To address this, we reconstructed 48 samples from four consecutive clock cycles (64 samples in total), aligned to a specific laser trigger onset (**Supplementary figure 2,** orange).

For example, suppose the laser trigger crosses the threshold at the 7th sample within the first set of 16 samples (①). In this case, the remaining 10 samples from A1, all 16 samples from the subsequent sets ② and ③, and the first 6 samples from ④ were combined to yield 48 samples, corresponding to 15 nanoseconds of data (denoted as ①', ②', ③' in orange). During this process, a second laser trigger also crossed the threshold. Accordingly, another set of 48 samples was concurrently processed starting from this second trigger event (③'', ④'', ⑤'' in green). While this second set was still being processed, a third laser trigger crossed the threshold. However, by this time, the processing of the first trigger had already completed, allowing the two circuits to alternate and handle triggers in parallel.

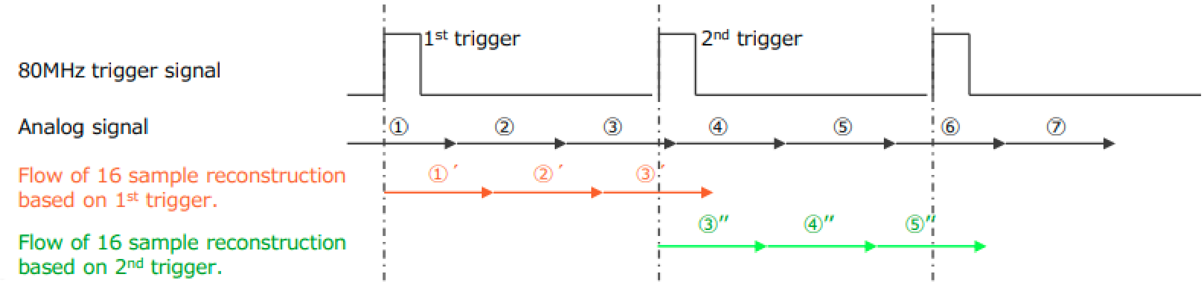


**Supplementary figure 2 | Timing diagram for reconstructing 48 analog samples synchronized to an 80 MHz trigger signal**

The top black trace shows the 80 MHz laser-trigger signal. The black arrows (① to ⑦) illustrates an example in which the analog signal is divided into three 5-ns sampling windows (16 samples at 3.2GS/s) per 80 MHz cycle. Because the analog signal reaches the system slightly after the laser trigger, reconstruction is offset accordingly. In the bottom trace (orange and green), three successive 16-sample segments are reconstructed for each trigger, giving 48 samples in total. Processing is parallelized for odd and even triggers, indicated in orange and green, so that a continuous stream of samples is maintained.

#### 2-4. Signal processing chain based on laser trigger and PMT signals.

In "trigger detection with hysteresis" (**Supplementary figure 3**) the laser trigger signal was detected at 3.2 GS/s and then counted in "gate for trigger”. If the number of lines was less than or equal to the number of lines in each frame, this was used as the trigger and the recording proceeded. This count was reset when the new V-Sync signal arrived. With "trigger distribution," the laser trigger was distributed from this single source to two circuit blocks (block A and block B) inside the single FPGA circuit.

The fluorescence intensity corresponding to each focus was detected at 80 MHz, but the pixel duel time in a typical two-photon microscope is longer than 12.5ns, which indicates that the single pixel signal includes fluorescence intensity for several excitation pulses. Therefore, in "averaging," we have made it possible for the FPGA to output the value for two or four laser pulses by averaging them. This successfully reduced the load on the hard disk. We configured up to four focal points for each laser trigger. Therefore, we set up four channels from 48 samples, and in each channel, we could determine which samples in the total 48 samples were summed up. The maximum number of samples to be summed was set to 16, and 12 bits of input were added to make 16 bits in the "summation". As a result, the FPGA outputs four 16-bit values per 1, 2 or 4 laser pulses. The PC transfer rate was 2 bytes (16 bit) x X channel x 80MHz trigger rate /Y average = 640*X/Y MB/s, where X was the number of channels and Y was the number of averaging. The maximum load was 640 MB/s when X=4, Y=1, and the minimum load was 40 MB/s when X=1, Y=4.

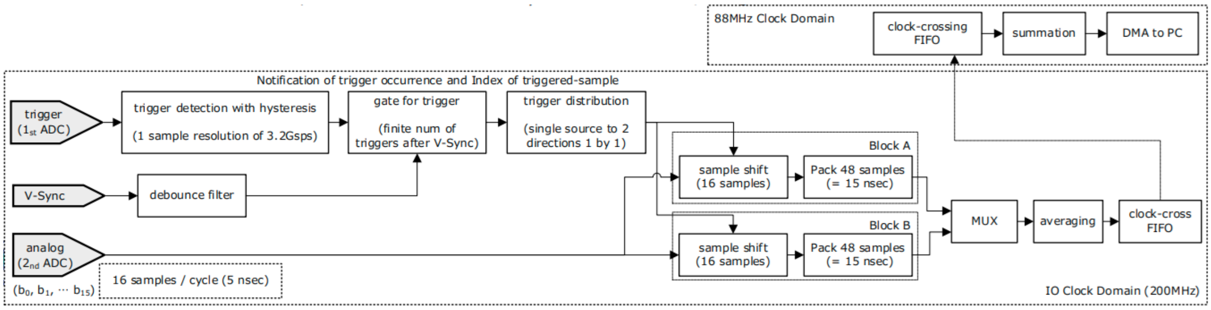


**Supplementary figure 3 | Signal processing chain**

The trigger signal acquired at 3.2 GS/s is detected with one-sample precision by a hysteresis discriminator. A V-Sync-synchronized gate accepts only valid triggers and reports their occurrence time together with the corresponding sample index. Each detected trigger is alternately routed to two paths, where the starting point of the analog data is shifted by 16 samples. Blocks A and B each reconstruct 48 samples, corresponding to a 15-ns window. Within the 200 MHz domain the two data streams are merged by a multiplexer and an averaging stage, then transferred to the 88 MHz domain through a clock-crossing FIFO. After passing through an adder the data are delivered to the PC in real time via DMA.

#### 3. Instruction of the LabVIEW software

The software has six libraries (sets of sub VIs). **Diesel2p Acquisition** directly controls the PXIe-5774 digitizer and its custom FPGA, handling PMT signal capture as well as hardware parameters such as trigger conditions and gain. The analog waveforms acquired are streamed at high-speed to the SSD by **TDMS Async Write-Analog**, while the corresponding line-synchronization signals are saved in parallel by **TDMS Async Write-H-Sync**. For offline analysis, **Diesel2p File Read** reconstructs and displays images from the stored analog and H-Sync data, and **Diesel2p File Conversion** batch-converts those images to TIFF format. **Raw acquisition** controls PXIe-5774 digitizer and its custom FPGA to record raw signals without compression. Together, these libraries provide a seamless path from data acquisition through visualization to file-format conversion.

Combining the five libraries, four top-level VIs implement the practical workflow. **tc-All Data Path for Display Data.vi** calls the Acquisition API to render real-time images for online monitoring. **All Data Path for Logging.vi** invokes the Acquisition API together with both TDMS Write APIs to save raw analog and digital data without loss. **Data Viewer.vi** employs the File Read API to replay previously acquired data as movies, whereas **File Conversion for Diesel2p.vi** uses the File Conversion API for exporting images as a tiff format. Separating these functions into dedicated VIs allows acquisition, storage, playback, and conversion to be executed and debugged independently.

Inside each VI, a hierarchical set of sub-VIs divides responsibilities finely. For example, sub-VIs in the Acquisition library abstract delay compensation for DI channels, trigger configuration, and integration-window settings so that they can be modified on-the-fly from the GUI. The “Context Help” window instantly reveals the purpose and default values of every parameter. This modular architecture enables researchers to create new VIs by re-using the base libraries and to extend the system with minimal code changes.

The following sections describe the VIs and libraries (sub-VIs) in detail. The same information is also available in each VI's help documentation.

### 3-1. VIs

- ➢ **tc-All Data Path for Display Data v3.vi**
- ➢ **All Data Path for Logging.vi**
- ➢ **Data Viewer v4.vi**
- ➢ **Read H-Sync and Analog TDMS and Write Multiple Image Files per Image.vi**

### 3-2. Libraries (Sets of sub-VIs):

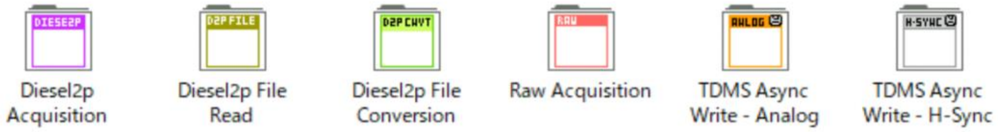


#### ➢ Diesel2p Acquisition

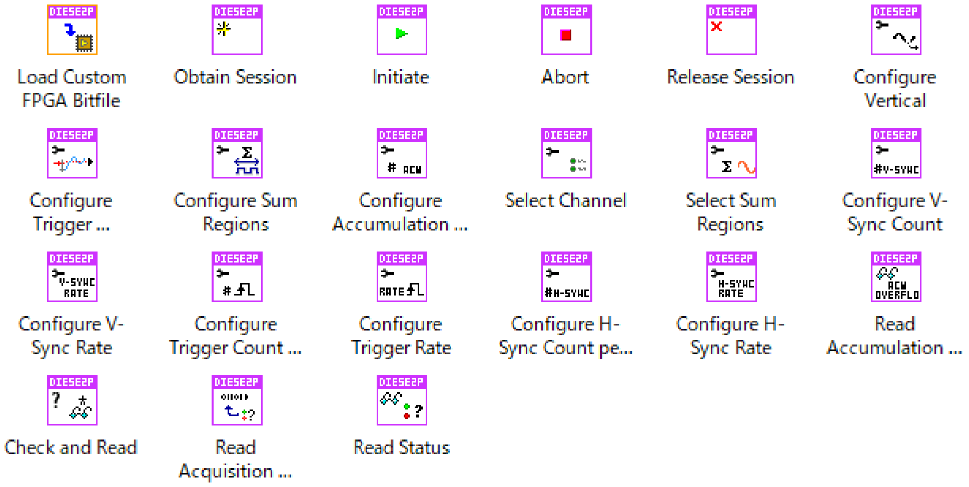


API to control the PXIe-5774 digitizer, a custom FPGA for FPGA-based Diesel2p measurements.

#### ➢ Diesel2p File Read

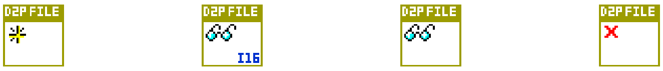

Obtain Session | Read One Image I16 | Read One Image | Release Session

API to reconstruct and display images based on stored analog and H-Sync signal data.

- **Diesel2p File Conversion**

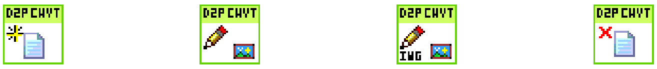

Obtain Session | Write 2D Array to Binary File | Write 2D Array to Image File | Release Session

API for reconstructing images from stored analog and H-Sync signal data and outputting them as TIFF-format image files.

- **Raw acquisition**

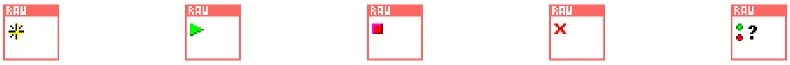

Obtain Session | Initiate | Abort | Release Session | Check Status

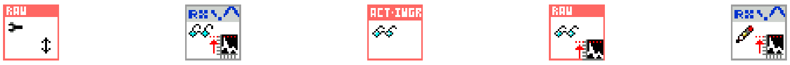

Configure Vertical | Read Scaling Info from TDM... | Read | Read and Scale | Write Scaling Info to TDMS ...

- **TDMS Async Write – Ananlog**

Diesel2p analog signal acquired by PXIe-5774 in combination with Diesel2p Acquisition's API Diesel2p analog signal acquired by 5774.

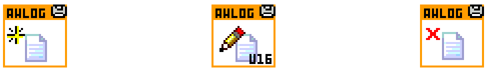

Obtain Session | Write Data - DVR - U16 | Release Session

- **TDMS Async Write – H-Sync**

API for high-speed storage of Diesel2p H-Sync data acquired by PXIe-5774 on an SSD on a PC in combination with Diesel2p Acquisition API.

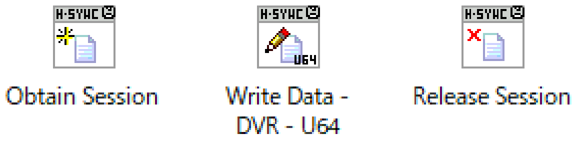


### 3-3. Description of each sub-VI:

➢ **Diesel2p Acquisition.lvclass:Load Custom FPGA Bitfile.vi**

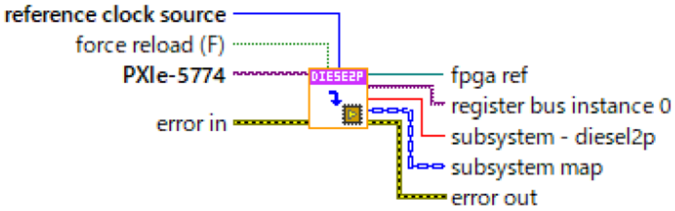


Load custom FPGA Bitfiles to PXIe-5774 device.

➢ **Diesel2p Acquisition.lvclass:Obtain Session.vi**

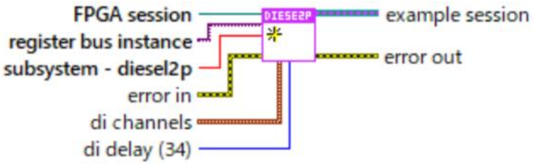


This function initializes a session of FPGA-based diesel2p measurements. Three references are required; **FPGA session**, **register bus instance**, and **subsystem - diesel2p**. These references are provided by **Load Custom FPGA Bitfile.vi**.

**di channels** specify Digital Input channels for V-Sync, H-Sync1, and H-Sync2. Default selections are DI2 for V-Sync, DI0 for H-Sync1, and DI1 for H-Sync2. The reason for the default selections is that SCB-12 terminal block is expected to be used. SCB-12 has two SMA inputs for DI0 and DI1, and the rest of the 6 digital inputs are spring terminals. Therefore, for solid connections of H-Sync, DI0 and DI1 are selected for H-Sync1 and 2, while DI2 is selected for V-Sync.

**di delay (34)** adjusts the delay of Digital Input data inside FPGA. ADC has pipeline delay and analog signals on FPGA arrive slightly after digital signals. To compensate for the time offset between the arrivals of analog and digital, digital signals are delayed inside the FPGA. For PXIe-5774, about 34 clock cycles delay gives a fairly good adjustment for the arrival offset.

FPGA-based diesel2p measurement is initialized with the following default configurations.

- trigger rate [Hz]: 80M
- trigger count: 1,000
- H-sync rate [Hz]: 8k
- H-sync count: 10
- sum region 1: enabled
- sum region 2: enabled
- sum region 3: enabled
- sum region 4: enabled
- sum region 1 start / length: 0/8
- sum region 2 start/ length: 0/8
- sum region 3 start / length: 0/8
- sum region 4 start/ length: 0/8
- vertical settings AI 0: 1Vpp, 0Voffset
- vertical settings AI 1: 1Vpp, 0Voffset
- accumulation count: 1
- channel select of data: AI 0 of PXIe-5774
- channel select of trigger: AI 1 of PXIe-5774

These configurations can be changed by other APIs of **Diesel2p Acquisition.**

➢ **Diesel2p Acquisition.lvclass:Intiate.vi**

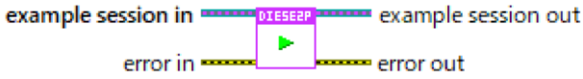


This function initiates Diesel2p acquisition.

➢ **Diesel2p Acquisition.lvclass:Abort.vi**

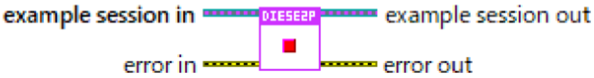

This function aborts Diesel2p acquisition.

➢ **Diesel2p Acquisition.lvclass:Release Session.vi**

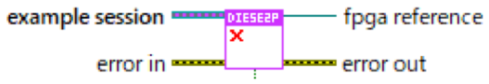


This function releases an obtained reference to the session.

➢ **Diesel2p Acquisition.lvclass:Configure Vertical.vi**

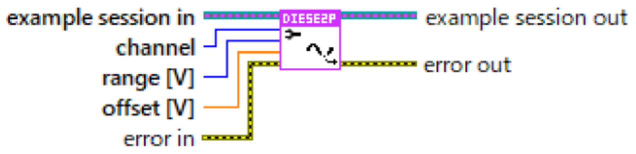


Based on the input configuration parameters, the gain, offset, etc. on the 5774 are configured via FPGA. It also internally calculates and stores the scale value for the binary data to be read according to the configuration.

**channels** specify the channels to be configured.

**range (Vpp)** specifies the input range.

**offset** specifies the offset.

➢ **Diesel2p Acquisition.lvclass:Configure Trigger Condition.vi**

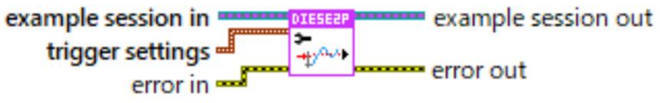


Configures trigger conditions to detect analog triggers for diesel2p measurement. the selected channel source with the specified trigger level, hysteresis percentage and trigger slope. You must run the **Select Channel VI** before running this VI.

**trigger level (V)** specifies the voltage threshold for the edge trigger.

**hysteresis level (%fs)** specifies the size of the trigger hysteresis window as a percentage of the configured vertical range. If the trigger slope is positive, the hysteresis window is below the trigger level. If the trigger slope is negative, the hysteresis window is above the trigger level. The digitizer triggers when the trigger signal crosses the trigger level after passing through the entire hysteresis window with the specified slope polarity.

**trigger slope** specifies either a positive slope or a negative slope for an edge trigger.

- **Diesel2p Acquisition.lvclass:Configure Sum Regions.vi**

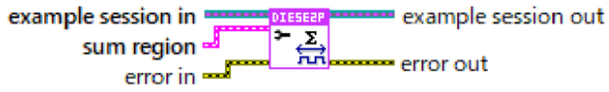


This function specifies the region in the 48-sample long record to sum up all the samples to get one scalar value. Arbitrary regions in 48 sample-long records can be specified by **start** and **length**. Max length is 16 samples. Four different regions can be selected, because FPGA-based diesel2p measurement allows to calculate multiple summations for different regions up to 4. The calculated sum value is 16 bit scalar output to support summation up to 16 12-bit samples (at most). Enable and disable multiple regions are configured by another VI named **Select Sum Regions.vi.**

- **Diesel2p Acquisition.lvclass:Configure Accumulation Count.vi**

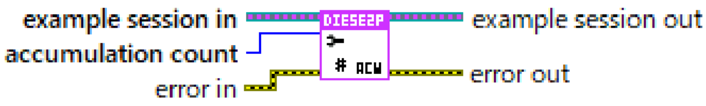


This function configures the count of accumulation for 48-samples long records. In a diesel2p system, analog trigger signals arrive at an 80MHz interval. a 48-sample long record is acquired at every trigger. When **accumulation count** is set to 1, no accumulation is applied. When the **accumulation count** is set to 2 or 4, 2 or 4 records are accumulated to a 14-bit internal cache. For example, when accumulation count is set to 2, accumulation is done after two triggers. After accumulation is finished, the accumulated record (48- sample long) is divided by the same number of accumulation counts to yield an average record.

- **Diesel2p Acquisition.lvclass:Select Channel.vi**

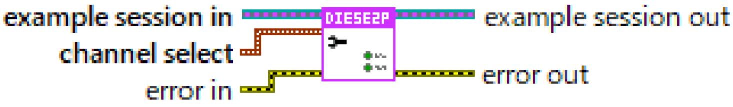


This function specifies assignments of two Analog Input channels on PXIe-5774. In diesel2p system, analog signal and analog trigger are the pair of analog signal source. **data** channel and **trigger** channel specify which of two analog input channels on PXIe-5774 is used to measure.

- ➢ **Diesel2p Acquisition.lvclass:Select Sum Regions.vi**

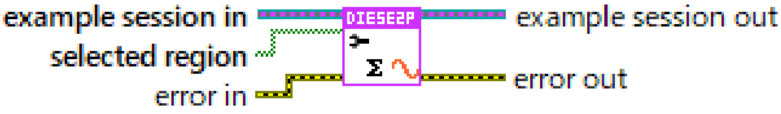


- ➢ **Diesel2p Acquisition.lvclass:Configure Trigger Count per V-Sync.vi**

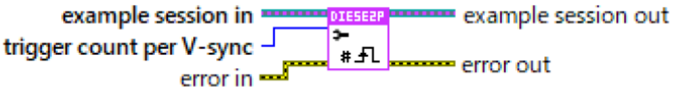


This function configures the number of analog triggers to detect after V-sync is received. If trigger count per V-sync is too many so that the next V-sync arrives during detecting triggers based on the previous V-sync, that next V-sync is ignored. Therefore, this function automatically checks if the configured trigger count per V-sync is too many so that next V-sync is ignored. At initialization of the FPGA-based diesel2p session, default trigger rate and trigger count are configured at 80MHz and 1,000.

- ➢ **Diesel2p Acquisition.lvclass:Configure Trigger Rate.vi**

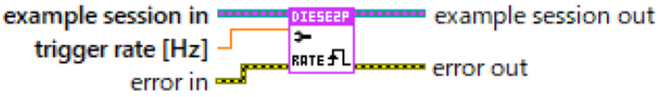


- ➢ **Diesel2p Acquisition.lvclass:Configure H-Sync Count per V-Sync.vi**

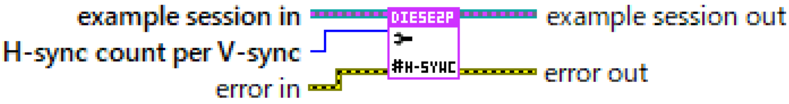

This function configures the number of H-sync to detect after V- sync is received. If H-sync count per V-sync is too many so that the next V-sync arrives during detecting H-sync based on the previous V-sync, the next V-sync is ignored. Therefore, this function automatically checks if the configured H- sync count per V-sync is too many. At initialization of the FPGA-based diesel2p session, default H-sync rate and H-sync count are configured 8kHz and 10.

- **Diesel2p Acquisition.lvclass:Configure H-Sync Rate.vi**

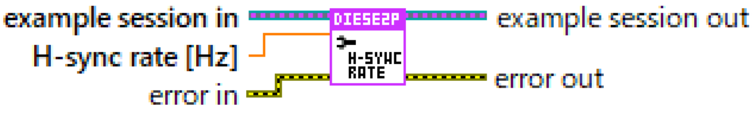


- **Diesel2p Acquisition.lvclass:Read Accumulation Overflow.vi**

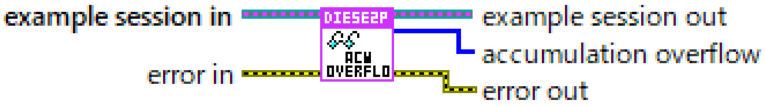


- **Diesel2p Acquisition.lvclass:Check and Read.vi**

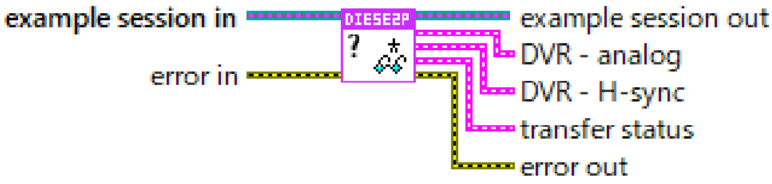


This function checks whether the host-side DMA buffer contains a specific number of samples, and if the number of samples in the DMA buffer exceeds that number, retrieves the same number of samples from the DMA buffer via Data Value Reference (DVR).

-the number of samples to check for the DMA buffer is calculated in advance, when acquisition is configured before it is started.

-the number of samples to check is calculated through acquisition configuration, so that data transfer rate from FPGA through host PC to external storage gets as efficient and fast as possible.

When samples are retrieved from the PC-side buffer, it is notified by TRUE output available along with the DVR. In addition, **read size** and **ch count** are returned to manipulate data, if that data is copied from DVR.

Above operations are executed for both analog data and for H-sync data. DVR, available, read size, ch count, and other output for the each of analog and H-sync are bundled into a cluster to output via single connector pane.

- **DVR - analog** for analog data
- **DVR - H-sync** for H-sync data

Data in analog data DVR is carried by samples of each enabled sum region in an interleaved format. For example, if sum regions of 0 and 2 are enabled out of four sum regions, summation results at 0 and 2 are interleaved. If trigger count per V-sync is configured to 1,000, 2,000 samples are returned in total, because 2 sum regions are enabled.

Refer to these two information clusters to separately handle analog data and H-sync data transferred from FPGA to PC.

➢ **Diesel2p Acquisition.lvclass:Read Acquisition State.vi**

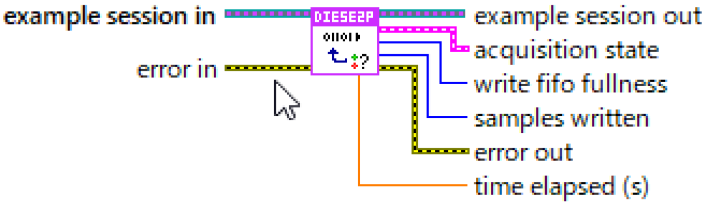


This function acquires the progress of data collection.

**acquisition state** acquires the acquisition state of the data acquisition engine on the FPGA.

**acquisition state.idle** is the data acquisition state in standby mode, waiting for the data collection instruction from the host PC side.

**acquisition state.in progress** is the state that data collection is active.

**acquisition state.write fifo**: overflow is the state that data transfer from FPGA to PC could not be completed in time and overflow occurred on the FPGA side.

**samples written** returns the number of samples transferred to the PC side since data acquisition started.

**time elapsed (s)** returns the time elapsed since data collection started.

**write fifo fullness (s)** returns the number of samples residing in the DRAM buffer in number of addresses.

➢ **Diesel2p Acquisition.lvclass:Read Status.vi**

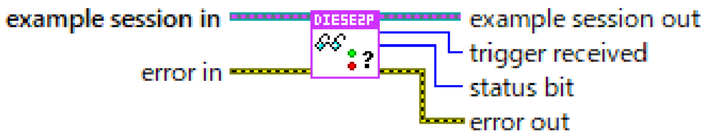


This function acquires the status information.

➢ **Diesel2p File Read.lvclass:Obtain Session.vi**

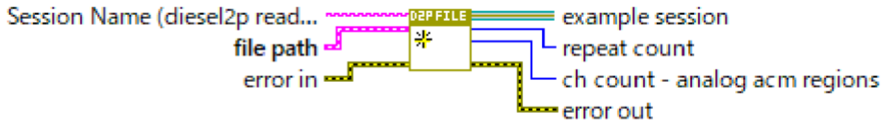


This function obtains a session.

➢ **Diesel2p File Read.lvclass:Read One Image I16.vi**

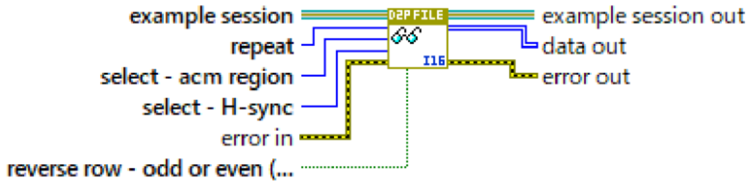


This function reads binary data for constructing an image.

➢ **Diesel2p File Read.lvclass:Release Session.vi**

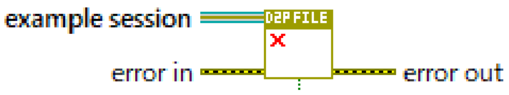


This function releases a obtained reference to the session.

➢ **Diesel2p File Conversion.lvclass:Obtain Session.vi**

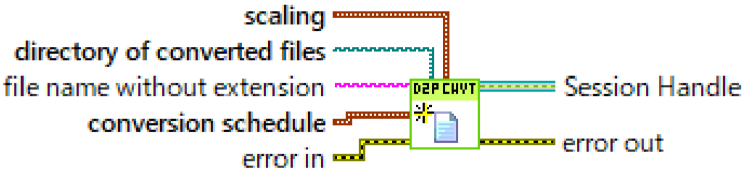


This function obtains a session for file conversion.

➢ **Diesel2p File Conversion.lvclass:Write 2D Array to Binary File.vi**

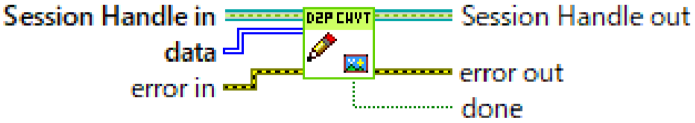


This function converts 2D arrays to binary data files.

➢ **Diesel2p File Conversion.lvclass:Write 2D Array to Image File.vi**

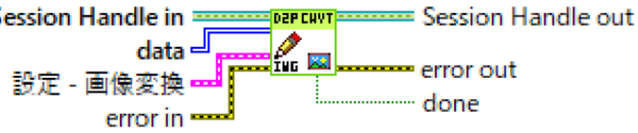


This function converts 2D arrays to image data files.

➢ **Diesel2p File Conversion.lvclass:Release Session.vi**

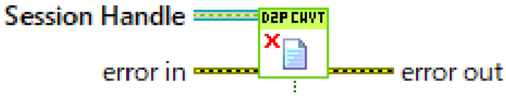


This function releases a obtained reference to the session.

➢ **Raw Acquisition.lvclass:Obtain Session.vi**

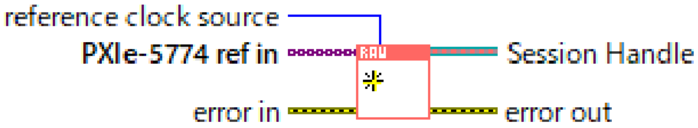


This function loads custom FPGA bitfiles to PXIe-5775 devices in the list.

For every PXIe-5775 device, wait until ADCs on PXIe-5775 settle, and configure the reference clock to PXI ClK10 of PXI chassis backplane. By configuring PXI_Clk10 as a reference clock for all devices, they refer to common clock signals by PLL to its IO clock.

Configure the master device to assert start trigger to all the other slave devices, when acquisition is started afterward by calling: Initiate (SubVI).vi. Then, synchronize all devices by TClk.

Configure default settings as follows.

1. acquisition length: 1000 samples
2. samples to read per channel: 1000 samples
3. decimation: : 1/8

➢ **Raw Acquisition.lvclass:Initiate.vi**

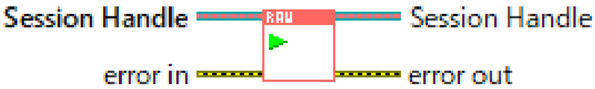


Start synchronized signal acquisition and processing. Start DMA FIFO and arm for start trigger on all listed PXIe-5775. Then, assert start to Master device which leads synchronized starts off all PXIe-5775 by TCIK. Then, the host side API transfers to the acquisition state.

➢ **Raw Acquisition.lvclass:Abort.vi**

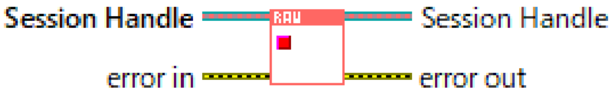


Abort acquisition and process. Reset and flush all data processes and data movement.

➢ **Raw Acquisition.lvclass:Release Session.vi**

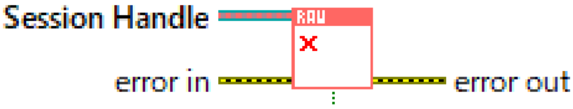


If a session is released during active operation, reset all operations and flush data from DMA bus buffers.

➢ **Raw Acquisition.lvclass:Check Status.vi**

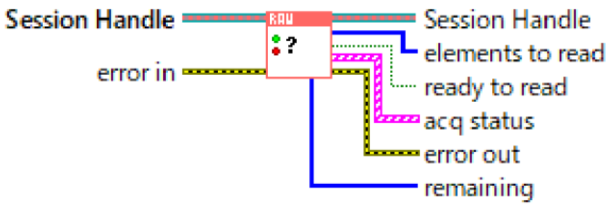


Check the number of samples transferred to the host PC and available inside the host-side DMA buffer. ready to read output notifies the timing to call Read API, once all the configured PXIe-5775 have transferred the number of samples which is configured by calling Set Number of Samples to Read per Channel.

➢ **Raw Acquisition.lvclass:Configure Vertical.vi**

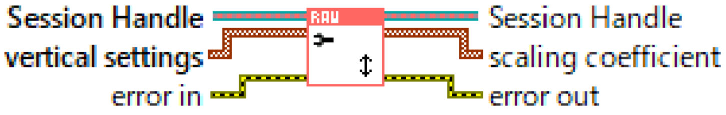


Start synchronized signal acquisition and processing. Start DMA FIFO and arm for start trigger on all listed PXIe-5775. Then, assert start to Master device which leads synchronized starts off all PXIe-5775 by TCIK. Then, the host side API transfers to the acquisition state.

➢ **Read Scaling Info from TDMS Property.vi**

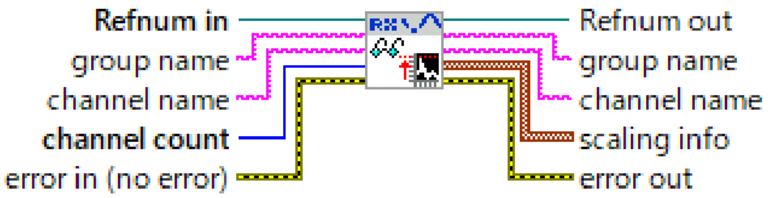


➢ **Raw Acquisition.lvclass:Read.vi**

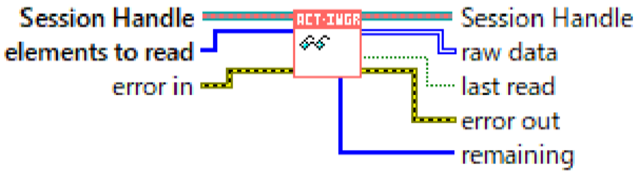


Read samples from host-side DMA FIFO of PXIe-5775 devices configured in session. elements to read specifies the number of samples to read from all devices. IMPORTANT NOTICE is elements to read should be retrieved from another API Check Status, which should be called right before every call of this API.

If scale is set FALSE, IQ data from FPGA is read in DBL format without scaling, which means FXP<+-,24,9> data of IQ channels are just converted into double-float precision format. On the other hand, if scale is set TRUE, data is scaled to voltage.

- **Raw Acquisition.lvclass:Read and Scale.vi**

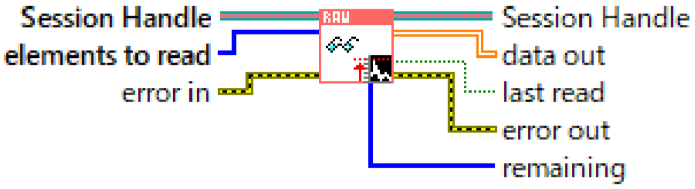


Read samples from host-side DMA FIFO of PXIe-5775 devices configured in session. elements to read specifies the number of samples to read from all devices. IMPORTANT NOTICE is elements to read should be retrieved from another API Check Status, which should be called right before every call of this API.

If scale is set FALSE, IQ data from FPGA is read in DBL format without scaling, which means FXP<+-,24,9> data of IQ channels are just converted into double-float precision format. On the other hand, if scale is set TRUE, data is scaled to voltage.

- **Raw Acquisition.lvclass:Write Scaling Info to TDMS Property.vi**

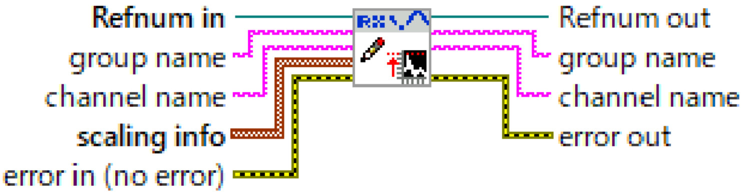


This function adds scaling info to TDMS property.

- **TDMS Async WRite - ANalong.lvclass:Obrain Session.vi**

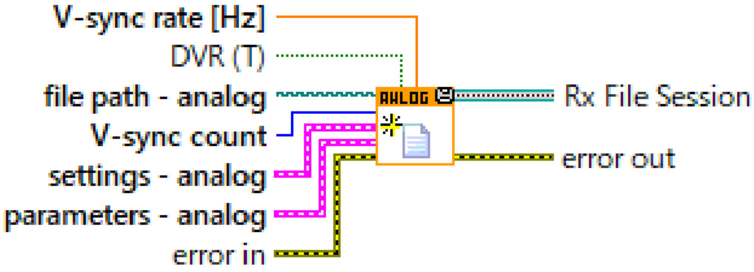


Initialize asynchronous TDMS file write operation for analog data acquired by FPGA-based Diesel2p system.

➢ **TDMS Async Write - ANalong.lvclass:Write Data -DVR - U16.vi**

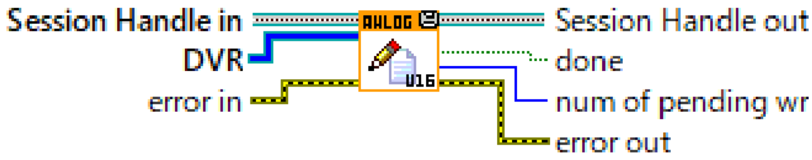


Write U16 data into TDMS file. DVR is Data Value Reference of data to be written to TDMS file. num of pending wr is number of data blocks waiting in queue for asynchronous file write operation.

➢ **TDMS Async Write - ANalong.lvclass:Release Session.vi**

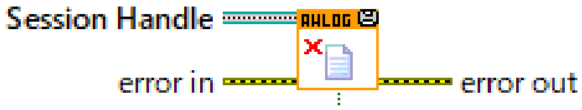


Release a obtained reference to the session.

➢ **TTDMS Async Write - H-Sync.lvclass:Obtain Session.vi**

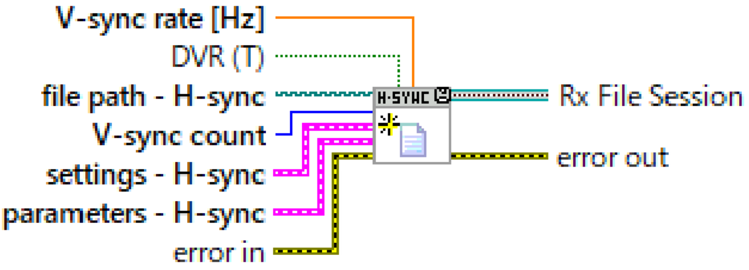


Initialize asynchronous TDMS file write operation for H-sync data acquired by FPGA-based Diesel2p system.

➢ **TTDMS Async Write - H-Sync.lvclass:Write Data -DVR - U64.vi**

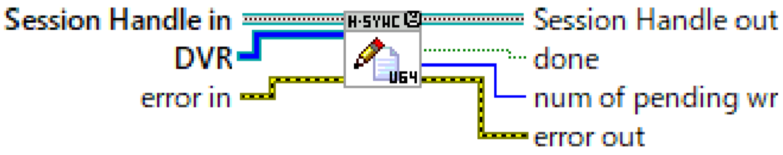


Write H-sync data into TDMS file. **DVR** is Data Value Reference of data to be written to a TDMS file. **num of pending wr** is number of data blocks waiting in queue for asynchronous file write operation.

➢ **TTDMS Async Write - H-Sync.lvclass:Release Session.vi**

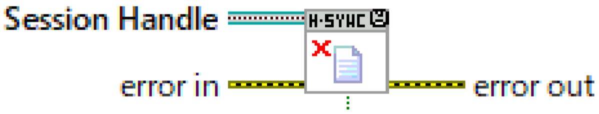


Release a obtained reference to the session.

**4. Modified optical parameters for Diesel2p**

Due to the difficulty in obtaining some glass materials during the global pandemic and other factors, as well as the rising prices of some glass materials, we attempted to keep as much of the original Diesel2p as possible while changing the lens configuration. The table (**Supplementary Figure 4**) details the optical design of our redesigned Diesel2p. Sub-assemblies 8-16, 18-26, and 34-42 are the same relay lenses. Sub-assembly 44-53 is a tube lens. Sub-assembly 56-69 is an objective lens. Mirrors 4 and 30 are galvo mirrors. The incidence light passes through the relay lens (LSM54-1050.ZBB, Thorlabs) and two concave lenses (AC508-250-B-ML, Thorlabs) after reflection from the resonant scanner (CRS8kHS, Cambridge) as previously described (Yu et al., supplementary figure 1). Several glasses highlighted in yellow were changed as follows, N-BK7 (517-642, refractive index) in the relay lens was replaced by S-BSL7 (516-641), and that in the tube lens was replaced by S-FPM3 (595-677). N-SF57 (847-238) was replaced with S-TIH53 (847-238). LF5 (581-410) was replaced with S-TIL25 (581-407). S-FPL51 (486-845) was replaced with S-FPL55 (486-701). N-SK2 (607-567) was replaced with S-BSM2 (607-568). Overall, the properties of the glass were almost the same as those of the original Diesel2p. Starting with the original ones, all optical parameters (thickness, radius, semi-diameter) were re-optimized with the Zemax software (OpticStudio). We confirmed that most optical features of this modified Diesel2p were comparable to those of the original one. Thus, we were able to build Diesel2p, which is less expensive and easier to manufacture, at least in Japan.

| # | Type | Comment | Radius | Thickness | Material | Semi-Diameter | Mech Semi-Diameter |
|---|---|---|---|---|---|---|---|
| 0 | STANDARD | | Infinity | Infinity | | 0.000 | 0.000 |
| 1 | STANDARD | | Infinity | 10.000 | | 4.050 | 4.050 |
| 2 | COORDBRK | | Infinity | 0.000 | | 0.000 | 0.000 |
| 3 | COORDBRK | | Infinity | 0.000 | | 0.000 | 0.000 |
| 4 | STANDARD | | Infinity | 0.000 | MIRROR | 7.371 | 7.371 |
| 5 | COORDBRK | | Infinity | 0.000 | | 0.000 | 0.000 |
| 6 | COORDBRK | | Infinity | -15.400 | | 0.000 | 0.000 |
| 7 | STANDARD | | Infinity | -2.000 | | 10.375 | 10.375 |
| 8 | STANDARD | Doublet 1 | 28.690 | -18.900 | S-PHM52 | 13.000 | 23.500 |
| 9 | STANDARD | | 15.831 | -11.700 | S-TIH53 | 15.000 | 23.500 |
| 10 | STANDARD | | 35.172 | -0.250 | | 23.500 | 23.500 |
| 11 | STANDARD | Doublet 2 | -92.300 | -14.100 | S-TIH53 | 30.000 | 30.000 |
| 12 | STANDARD | | -36.360 | -25.000 | S-LAH53 | 30.000 | 30.000 |
| 13 | STANDARD | | 73.760 | -39.650 | | 30.000 | 30.000 |
| 14 | STANDARD | Doublet 3 | 29.250 | -15.800 | S-TIH53 | 25.000 | 30.000 |
| 15 | STANDARD | | 36.190 | -9.400 | S-BSL7 | 30.000 | 30.000 |
| 16 | STANDARD | | 48.000 | -10.240 | | 30.000 | 30.000 |
| 17 | STANDARD | | Infinity | -10.240 | | 24.721 | 24.721 |
| 18 | STANDARD | | -48.000 | -9.400 | S-BSL7 | 30.000 | 30.000 |
| 19 | STANDARD | | -36.190 | -15.800 | S-TIH53 | 30.000 | 30.000 |
| 20 | STANDARD | | -29.250 | -39.650 | | 25.000 | 30.000 |
| 21 | STANDARD | | -73.760 | -25.000 | S-LAH53 | 30.000 | 30.000 |
| 22 | STANDARD | | 36.360 | -14.100 | S-TIH53 | 30.000 | 30.000 |
| 23 | STANDARD | | 92.300 | -0.250 | | 30.000 | 30.000 |
| 24 | STANDARD | | -35.172 | -11.700 | S-TIH53 | 23.500 | 23.500 |
| 25 | STANDARD | | -15.831 | -18.900 | S-PHM52 | 15.000 | 23.500 |
| 26 | STANDARD | | -28.690 | -2.000 | | 13.000 | 23.500 |
| 27 | STANDARD | | Infinity | -15.400 | | 10.375 | 10.375 |
| 28 | COORDBRK | | Infinity | 0.000 | | 0.000 | 0.000 |
| 29 | COORDBRK | | Infinity | 0.000 | | 0.000 | 0.000 |
| 30 | STANDARD | | Infinity | 0.000 | MIRROR | 6.363 | 6.363 |
| 31 | COORDBRK | | Infinity | 0.000 | | 0.000 | 0.000 |
| 32 | COORDBRK | | Infinity | 15.400 | | 0.000 | 0.000 |
| 33 | STANDARD | | Infinity | 2.000 | | 10.375 | 10.375 |
| 34 | STANDARD | | -28.690 | 18.900 | S-PHM52 | 13.000 | 23.500 |
| 35 | STANDARD | | -15.831 | 11.700 | S-TIH53 | 15.000 | 23.500 |
| 36 | STANDARD | | -35.172 | 0.250 | | 23.500 | 23.500 |
| 37 | STANDARD | | 92.300 | 14.100 | S-TIH53 | 30.000 | 30.000 |
| 38 | STANDARD | | 36.360 | 25.000 | S-LAH53 | 30.000 | 30.000 |
| 39 | STANDARD | | -73.760 | 39.650 | | 30.000 | 30.000 |
| 40 | STANDARD | | -29.250 | 15.800 | S-TIH53 | 25.000 | 30.000 |
| 41 | STANDARD | | -36.190 | 9.400 | S-BSL7 | 30.000 | 30.000 |
| 42 | STANDARD | | -48.000 | 10.240 | | 30.000 | 30.000 |
| 43 | STANDARD | | Infinity | 20.000 | | 24.829 | 24.829 |
| 44 | STANDARD | | Infinity | 75.000 | N-BK7 | 37.500 | 37.500 |
| 45 | STANDARD | | Infinity | 112.730 | | 37.500 | 37.500 |
| 46 | STANDARD | Singlet 1 | -54.893 | 19.800 | S-FPM3 | 42.500 | 50.000 |
| 47 | STANDARD | | -91.910 | 0.920 | | 50.000 | 50.000 |
| 48 | STANDARD | Doublet 4 | 575.210 | 25.000 | S-PHM52 | 50.000 | 50.000 |
| 49 | STANDARD | | -92.300 | 11.400 | S-TIL25 | 50.000 | 50.000 |
| 50 | STANDARD | | -133.200 | 119.970 | | 50.000 | 50.000 |
| 51 | STANDARD | Doublet 5 | 100.070 | 5.000 | S-TIL25 | 45.000 | 45.000 |
| 52 | STANDARD | | 56.000 | 24.800 | S-FPL55 | 45.000 | 45.000 |
| 53 | STANDARD | | 141.567 | 157.060 | | 45.000 | 45.000 |
| 54 | STANDARD | | Infinity | 0.000 | | 18.440 | 18.440 |
| 55 | STANDARD | Obj | Infinity | -13.530 | | 16.962 | 16.962 |
| 56 | STANDARD | | -39.770 | 14.700 | S-TIH6 | 22.000 | 26.750 |
| 57 | STANDARD | | 114.397 | 11.309 | | 26.750 | 26.750 |
| 58 | STANDARD | | 314.910 | 15.800 | S-BSM2 | 35.750 | 35.750 |
| 59 | STANDARD | | -69.390 | 29.340 | | 35.750 | 35.750 |
| 60 | STANDARD | | 320.000 | 25.000 | S-BSM2 | 45.000 | 45.000 |
| 61 | STANDARD | | -86.020 | 1.300 | | 45.000 | 45.000 |
| 62 | STANDARD | | 117.057 | 25.000 | S-PHM52 | 42.000 | 42.000 |
| 63 | STANDARD | | -65.260 | 7.200 | S-TIH53 | 42.000 | 42.000 |
| 64 | STANDARD | | 325.170 | 7.680 | | 42.000 | 42.000 |
| 65 | STANDARD | | Infinity | 0.000 | | 31.066 | 31.066 |
| 66 | STANDARD | | 48.960 | 20.400 | S-PHM52 | 35.600 | 35.600 |
| 67 | STANDARD | | 118.870 | 17.125 | | 30.200 | 35.600 |
| 68 | STANDARD | | 37.570 | 23.300 | S-TIH53 | 21.100 | 21.100 |
| 69 | STANDARD | | 29.723 | 1.733 | | 10.000 | 21.100 |
| 70 | STANDARD | | Infinity | 8.000 | | 8.283 | 8.283 |
| 71 | STANDARD | | Infinity | 0.000 | | 3.277 | 3.277 |
| 72 | STANDARD | Coverslip | Infinity | 0.150 | BK7 | 3.277 | 3.277 |
| 73 | STANDARD | Tissue | Infinity | 0.500 | SEAWATER | 3.220 | 3.277 |
| 74 | STANDARD | | Infinity | 0.000 | SEAWATER | 3.125 | 3.277 |

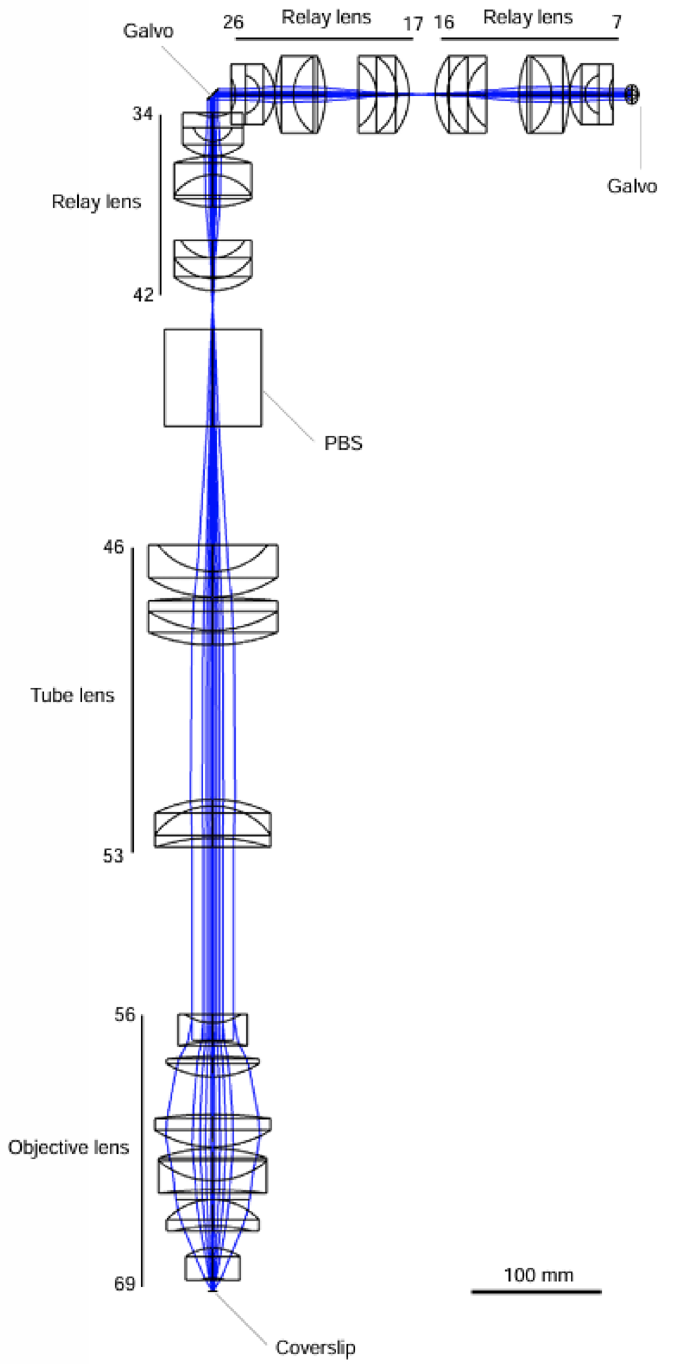

**Supplementary figure 4 | Optical design of the modified Diesel2p mesoscope.**

Left: Zemax lens-data editor listing all 74 sequential surfaces in the excitation path. Surfaces whose glass type was changed from the original Diesel2p design are highlighted in yellow. Columns indicate surface type, comment (e.g., singlet, doublet), radius of curvature, center thickness, glass material, and mechanical semi-diameter. Right: Paraxial ray trace of the excitation beam. A pair of galvanometric scanners directs the beam through three relay-lens groups (surfaces 7–41) to a polarizing beam-splitter (PBS, surface 42). The beam then passes a four-element tube lens (surfaces 46–53) and a seven-element objective lens (surfaces 56–69) before reaching the coverslip above the specimen. Numbers correspond to the sequential surface indices in the lens table; brackets denote the functional sub-assemblies.
